# Overexpression profiling reveals cellular requirements in context of genetic backgrounds and environments

**DOI:** 10.1101/2022.07.29.502095

**Authors:** Nozomu Saeki, Chie Yamamoto, Yuichi Eguchi, Takayuki Sekito, Shuji Shigenobu, Mami Yoshimura, Yoko Yashiroda, Charles Boone, Hisao Moriya

**Author notes:** **Author contribution:** SN and HK performed experiments, SN, HK, SS, TM, and HM analyzed the data, SN, HK, and HM prepared the figures and wrote the manuscript text. CB and SS supervised high-throughput sequencing analyses.

## Abstract

Overexpression due to copy number variation, promoter mutation, or aneuploidy is often observed, but its adaptive role is not clearly understood. Using a novel “overexpression profiling” method designated ADOPT, we systematically obtained genes whose overexpression was functionally adaptive (GOFAs) under stress conditions in budding yeast to elucidate the nature of adaptive overexpression. GOFAs obtained under heat, salt, and oxidative stress were unique genes that differed from known stress response genes. GOFAs under salt (NaCl) stress were genes involved in calcium homeostasis, reflecting the calcium deficiency of the medium. GOFAs from different genetic backgrounds and co-overexpressing strains revealed that calcium and potassium requirements in salt stress tolerance differ among strains, which is reflected. Profiling of the knockout collection suggested that the effect of calcium was to prevent mitochondrial outbursts. Mitochondria-enhancing GOFAs were adaptive only when calcium was sufficient and conversely non-adaptive in calcium deficiency, supporting the above hypothesis. Adaptive overexpression, thus, reflects the cellular requirements for maximizing the organism’s adaptive capacity within a given environmental and genetic context.

## Introduction

Organisms are able to sustain growth, reproduction, and survival even under stressful environmental conditions ^1^. This ability is achieved through evolutionarily-conserved intracellular systems such as signaling pathways and stress-induced proteins that play essential roles under stress ^2–4^. For example, a protein kinase complex mTOR is known to be an evolutionarily conserved hub of signaling pathways for cell proliferation and stress responses ^5^. Heat shock transcription factors are known to precisely regulate the expression of heat shock proteins, which are chaperone proteins and protect cell function ^6^. Thus, cell behavior in response to environmental stresses should be programmed and optimized ^7,8^. On the other hand, accidental increases in gene expression (i.e., overexpression) due to mutations such as single nucleotide polymorphisms (SNPs) and copy number variations (CNVs) are often observed in natural isolates. They are thought to enhance organismal adaptation under conditions of uncertainty ^9–12^.

However, it is poorly understood why and how overexpression can be adaptive despite well-established stress response systems. An exceptional but well-studied example of adaptive overexpression is drug resistance. Cells and individuals acquire drug resistance when specific genes are overexpressed ^13–16^. Those genes encode efflux pumps, drug-degrading enzymes, and drug targets that interact directly with drugs. In contrast to drug resistance, the effects of overexpression on environmental stresses cannot be readily assumed because of the combined effects of many factors.

In this study, using the model eukaryote *Saccharomyces cerevisiae* and the newly developed experimental system “ADOPT”, we systematically isolated <u>g</u>enes whose <u>o</u>verexpression is <u>f</u>unctionally <u>a</u>daptive (GOFAs) to understand their contribution to overcoming stress environments. GOFAs isolated under high temperature, high salt, and oxidative stress suggested that gene overexpression can compensate for deficiencies to maximize fitness under stress. The results also suggested that whether gene overexpression is adaptive or not is highly dependent on genetic background and environmental factors. Thus, it is possible to determine the missing piece for cell growth in each genetic background and environment by examining GOFAs.

## Results

### GOFAs under stress are a unique set of genes

To systematically identify GOFA, we set up an experimental system designated ADOPT (based on “<u>a</u>utonomous <u>d</u>osage-<u>o</u>ptimization using <u>p</u>lasmids with two-micron origin”), as shown in **Fig 1A**. The ADOPT system consists of four steps; 1) Construction of libraries pooling *S. cerevisiae* BY4741 cells that harbor a 2µ plasmid with each of the cloned 5,751 genes of the BY4741 genome. The initial version used two sets of gTOW6000 libraries ^17^ to build Pool_a and Pool_b; 2) Competitive culture and passaging of the pooled libraries; 3) Long-read plasmid sequencing of plasmid inserts extracted from the culture pool library by the Oxford Nanopore Technologies MinION **(Extended Data Fig. 1A**); 4) Analysis of sequence data to calculate the frequencies of plasmid inserts and identification of GOFAs. We confirmed that both Pool_a and Pool_b covered more than 93% of the 5,751 genes (**Extended Data Fig. 1B**).

**Figure 1.**
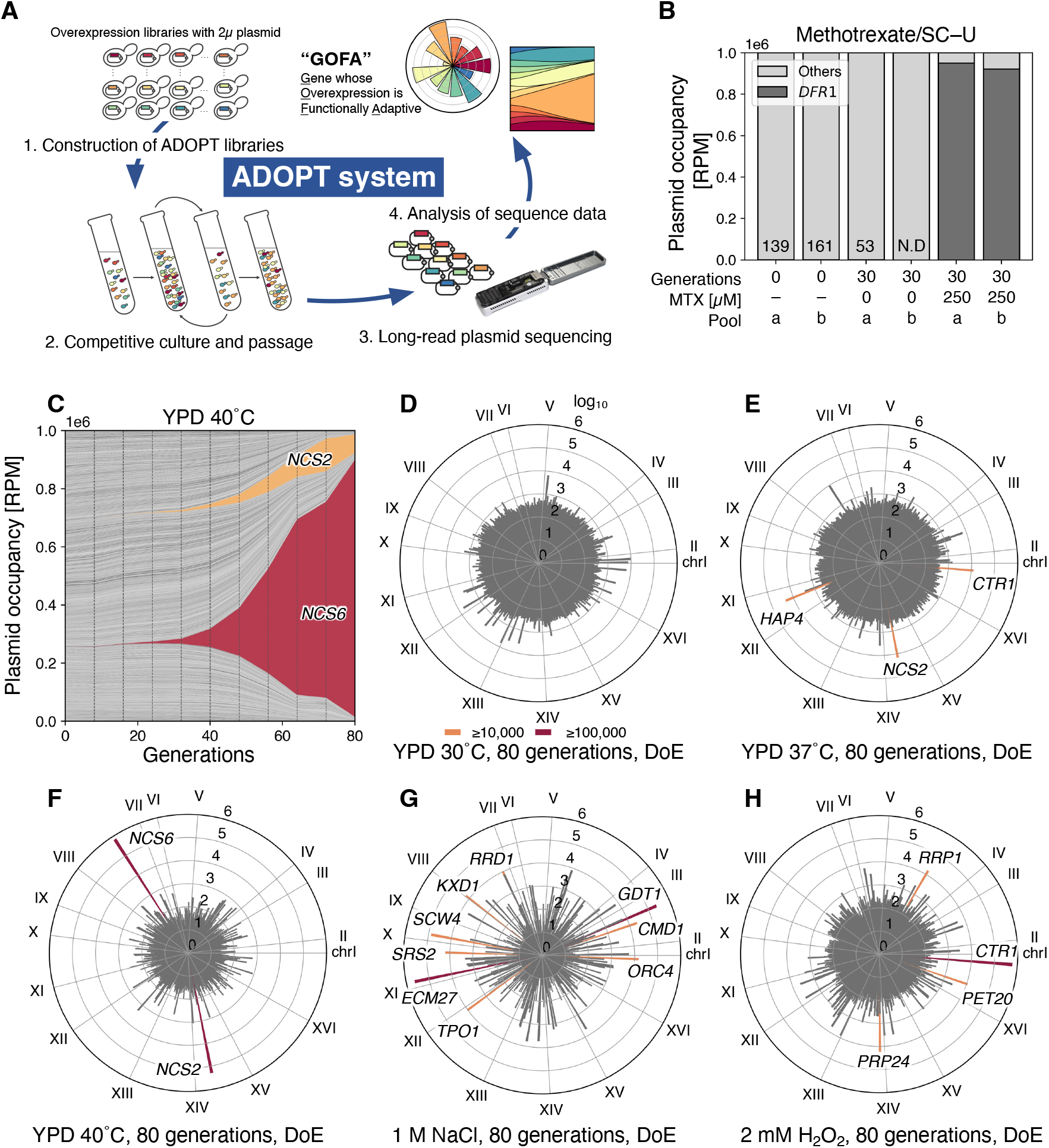
GOFAs under well-studied stresses are a unique set of genes. **A)** The ADOPT system for identifying GOFAs developed in this study. The detail is explained in the text. **B)** A proof of concept for ADOPT: identification of GOFA as a drug target (under 250 µM methotrexate). The bar plot and the numbers show occupancies of a plasmid harboring *DFR1* showing with reads per million (RPM) reads. **C)** An example of the result of the ADOPT experiment (under heat stress). One of the four replicates at 40°C in YPD for 80 generations (samples were analyzed every eight generations). Occupancies of each plasmid are shown with reads per million (RPM) reads, and the orange and red areas correspond to the enriched genes *NCS2* and *NCS6*, respectively. **D-H)** Degree of enrichment (DoE) of plasmid inserts after the cultivation of pooled libraries under well-studied stresses; YPD at 30°C (C, control), 37°C (D) and 40°C (E) as the heat stresses, 1 M NaCl (F) as the salt stress, 2 mM H2O2 (G) as the oxidative stress. Each cycle chart shows the average DoE of 5,751 genes in four replicates after 80 generations. Each gene is plotted with its chromosomal location (chr I to XVI). The gene names of GOFAs with DoE ≥ 100,000 (red bars) and DoE ≥ 10,000 (orange bars) are also shown. These data are summarized in Table S2.

As proof of concept for ADOPT, we screened for GOFA in the presence of 250 µM methotrexate (MTX)^18^, an antagonistic inhibitor of dihydrofolate reductase in yeast encoded by *DFR1*. As expected, 30 generations of competitive culture enriched the *DFR1* plasmid with more than 90% of the total (**Fig. 1B**). Next, using the ADOPT system, we identified GOFAs under the most studied stresses in yeast (**Extended Data Fig. 2A**), namely heat stress (37 °C and 40 °C), salt/osmotic stress (1 M NaCl) and oxidative stress (2 mM H_2_O_2_). The exact temperature and concentration of chemicals were determined to stress normal yeast growth (**Extended Data Fig. 2B-D**). **Fig. 1C** shows the time course of ADOPT analysis at 40°C. *NCS2* and *NCS6* were found to be dramatically enriched during incubation. Enriched genes were highly reproducible across replicates (**Extended Data Fig. 1D**), although slight differences were observed for different pools of origin (**Extended Data Fig. 1D-J)**.

There was a negative correlation between insert length and the number of reads, suggesting some technical bias in reading inserts (**Extended Data Fig. 3**). To reduce these analysis biases, we calculated the degree of enrichment (DoE, see **Method**) and defined genes with DoE higher than 10,000 as GOFA. **Fig. 1D-H** show the DoE of 5,751 genes in each condition. In YPD medium at 30°C, no genes were enriched even after 80 generations (**Fig. 1C**), suggesting that GOFAs were absent in this non-stressed condition. Moreover, most dose-sensitive genes ^17^ were shed from the culture as expected (**Extended Data Fig. 4A-C**) ^20^. At 37°C, *NCS2*, *CTR1*, and *HAP4*; at 40°C, *NCS2* and *NCS6* were enriched (**Fig. 1E and 1F**); under 1 M NaCl, nine genes, including *ECM27* and *GDT1*, were enriched; under H_2_O_2_, four genes, including *CTR1* (**Fig. 1G and 1H**) were enriched (**Fig. 1G and 1H**).

We initially expected that our ADOPT experiments would identify so-called “stress-responsive genes”. For example, overexpression of some heat shock-responsible genes might confer high-temperature tolerance. However, no above-mentioned GOFAs appear to be included in such categories. In fact, none of the enriched GOFAs matched stress-inducible genes such as “environmental stress-responsive genes” or “heat shock responsive genes” ^7,20^. The original libraries contained those genes, but they were excluded during the ADOPT experiments (**Extended Data Fig. 4**). These results suggested that GOFAs identified by our ADOPT system are a unique set of genes that might be helpful for an understanding of unexplored cellular physiology under stress.

To obtain clues to the nature of the identified GOFAs, we first focused on *NCS2* and *NCS6* at 40°C (**Fig. 1C and 1F**). Overexpression of *NCS2* and *NCS6* (*-oe*) independently resulted in increased heat resistance (**Extended Data Fig. 2E and 2F**). *NCS2* and *NCS6* are involved in the thiolation of wobble uridine of tRNAs in the *URM1* pathway ^21^, and the *URM1* pathway is known to have strain-dependent thermo-sensitivity ^22,23^. Since the BY4741 strain derived from S288C has a thermosensitive *URM1* pathway ^23^, we thought that *NCS2-oe* and *NCS6-oe* should compensate for their functions at high temperatures. Thus, we hypothesized that GOFAs generally function to compensate for cellular requirements in each environment.

### GOFAs enriched under salt stress propose Ca^2+^ limitation of the culture medium

To further ascertain the above hypothesis, we next focused on GOFAs enriched under salt stress (1 M NaCl, **Fig. 1G**). GOFAs enriched in all four replicates (*CMD1*, *GDT1*, and *ECM27*, **Fig. 2A**) were involved in intracellular calcium homeostasis (**Fig. 2B**); *CMD1* encodes calmodulin ^25,26^, while *GDT1* and *ECM27* encode calcium transporters that localize to the Golgi and ER membranes, respectively ^25,26^. *YBR196C-A*, enriched in Pool_b-derived replicates (**Fig. 2A**), has been reported as an “emerging gene” and encodes an adaptive protein that localizes to the ER membrane (**Fig. 2B**) ^27^. We confirmed that their overexpression significantly increased the growth rate under 1 M NaCl (**Fig. 2C**). We also confirmed the enhancement of the “calcium pulse” (rapid increases in cytosolic Ca^2+^ concentration) upon salt/osmotic stress exposure under *GDT1-oe* and *ECM27-oe* (**Fig. 2D**)^25,26,28,29^. *YBR196C-A-oe* also enhanced the calcium pulse even stronger (**Fig. 2D**). Based on these results and our above hypothesis; we speculated that GOFA under high salt might compensate for the calcium requirement under our experimental conditions. Indeed, it was true, as the addition of Ca^2+^ (5 and 20 mM) to the medium increased the growth rate of cells under salt stress (**Fig. 2E**). Furthermore, the increase in growth rate with Ca^2+^ addition canceled out the advantage of GOFAs under salt stress (**Fig. 2F**), suggesting that GOFAs mimic Ca^2+^ addition.

**Figure 2.**
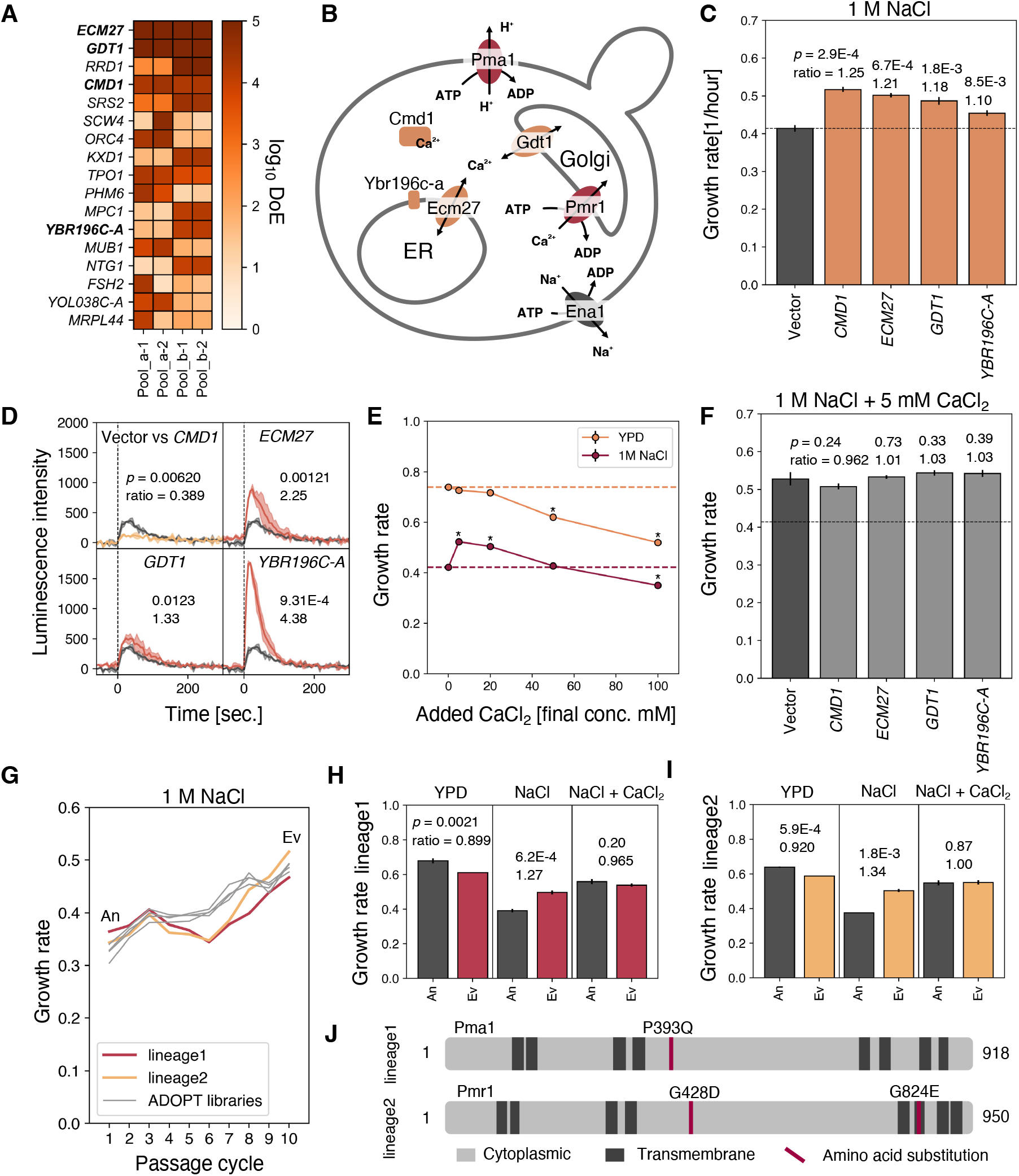
GOFAs as well as adaptive mutants under salt stress, propose Ca^2+^ limitation in the culture medium. **A)** Genes whose DoE was ≥10,000 in at least one of four replicates under the salt stress. Their DoE scores (log_10_ DoE) are shown with the darkness of the orange color. **B)** A cellular diagram illustrating protein functions in calcium homeostasis identified in this study. **C)** Growth rates of the cells overexpressing GOFAs under 1 M NaCl. **D)** The cytoplasmic Ca^2+^ pluses of the cells overexpressing GOFAs upon the salt stress measured by the aequorin luminescence assay. Grey and orange lines show the empty vector as control and targets, respectively. The vertical dashed lines represent the timing of added NaCl. **E)** The effect of CaCl_2_ addition on growth rates of BY4741 with (red) or without 1 M NaCl (orange). The horizontal dashed lines show the growth rate of BY4741 in YPD (orange dash) or 1 M NaCl without adding CaCl_2_ (red dash). Asterisks indicate significant differences. **F)** Growth rates of the cells overexpressing GOFAs under 1 M NaCl with 5 mM CaCl_2_. The horizontal dashed line presents the empty vector control’s growth rate without CaCl2 addition (shown in C). **G)** Changes in ADOPT pooled library growth rates and empty vector controls during the passages under salt stress. The linage1 (red) and linege2 (orange) of the vector controls and four replicates of ADOPT pooled library (grey) under 1 M NaCl are shown. For the vector control, the cells first inoculated were designated as “ancestor (An)” and the cells obtained after the 10th passage cycle as “evolved (Ev)”. **H and I)** Growth rates of An and Ev cells of linege1 (H) and linege2 (I) under YPD (left), 1 M NaCl (middle), and 1 M NaCl with 5 mM CaCl2 (right). **J)** Diagrams showing the amino acid substitutions in Pma1 and Pmr1 from the Ev lineage 1 and lineage 2. The dark grey areas indicate the transmembrane domains, and red bars indicate amino acid substitutions. The *p*-values are from Welch’s t-test (n = 3). The significance was evaluated by the Bonferroni correction (*p* < 0.05/4 = 0.0125). Error bars or the filled areas indicate SD.

During the ADOPT experiment under salt stress, we unexpectedly observed that the control strain without libraries also adapted (or “evolved” to adapt) to the salt stress (lineage1 and lineage2 in **Fig. 2G**). The two evolved lineages after 10 passages (Ev) grew significantly faster than the ancestral strain (An) under 1 M NaCl (**Fig. 2H and 2I**). We re-sequenced the entire genome of the Ev lineages and found base substitutions in the genome that cause P393 to Q in Pma1, G428 to D, and G824 to E in Pmr1 (**Fig. 2J**). Since Pma1 and Pmr1 are also involved in calcium homeostasis ^30,31^ (**Fig. 2B**), we speculated that cells may have also evolved to compensate for calcium deficiency under our conditions through these mutations. This idea also seemed correct since the increased growth rate due to the addition of Ca^2+^ under salt stress canceled out the advantage of the Ev strain (**Fig. 2H and 2I, right**).

The results so far indicate that the growth conditions used here lack sufficient Ca^2+^ for maximum salt tolerance of the yeast cells, resulting in the isolation of genes that may compensate for the Ca^2+^ requirement by overexpression (or mutation).

### GOFAs reflect differences in yeast strains

Previous studies have identified several genes confer salt tolerance through copy number variance (CNV) or overexpression. For example, CNV of *ENA1*, a sodium exporter, affects salt tolerance ^10,32^, and overexpression of *HAL1-9,* named HALotolerance, increases salt tolerance ^33–37^. We were thus surprised that the GOFAs identified in the 1 M NaCl conditions (**Fig. 1G and 2A**) did not contain those genes, even though they were present in our original libraries (**Table S3**). Because *S. cerevisiae* varies greatly in salt tolerance among strains ^10^ (**Extended Data Fig. 5A**), we speculated that strain differences might reflect GOFAs.

To test this possibility, we analyzed the effect of Ca^2+^ on the salt tolerance of different *S. cerevisiae strains*; laboratory strains BY4741 (a derivative of S288C used so far), W303, and CEN.PK2-1C (CEN.PK) and a European wine strain, DBVP6765 (**Fig. 3A**). We also analyzed several other strains, as shown in **Extended Data Fig. 5B**. As reported, their salt tolerance without Ca^2+^ was quite different; under 1 M NaCl, DBVP6765 grew much slower than BY4741 and W303, and CEN.PK did not grow (**Fig. 3A**, 0 mM Ca^2+^). Interestingly, adding Ca^2+^ dramatically improved the salt tolerance of DBVP6765 and CEN.PK; adding 5 mM Ca^2+^ nearly canceled the salt sensitivity of DBVP6765 compared to BY4741 and W303 (**Fig. 3A**) while adding up to 50 mM Ca^2+^ salt tolerance of CEN.PK increased gradually but significantly (**Fig. 3A**). Note that the addition of Ca^2+^ did not increase the growth rate of the strains without salt stress but decreased it (**Extended Data Fig. 5C-E**). These results suggest that the differences in salt sensitivity of each strain may be explained by differences in Ca^2+^ requirements, which could potentially reflect differences in GOFAs. Therefore, we next attempted to identify the GOFAs of CEN.PK and DBVP6765 strains.

**Figure 3.**
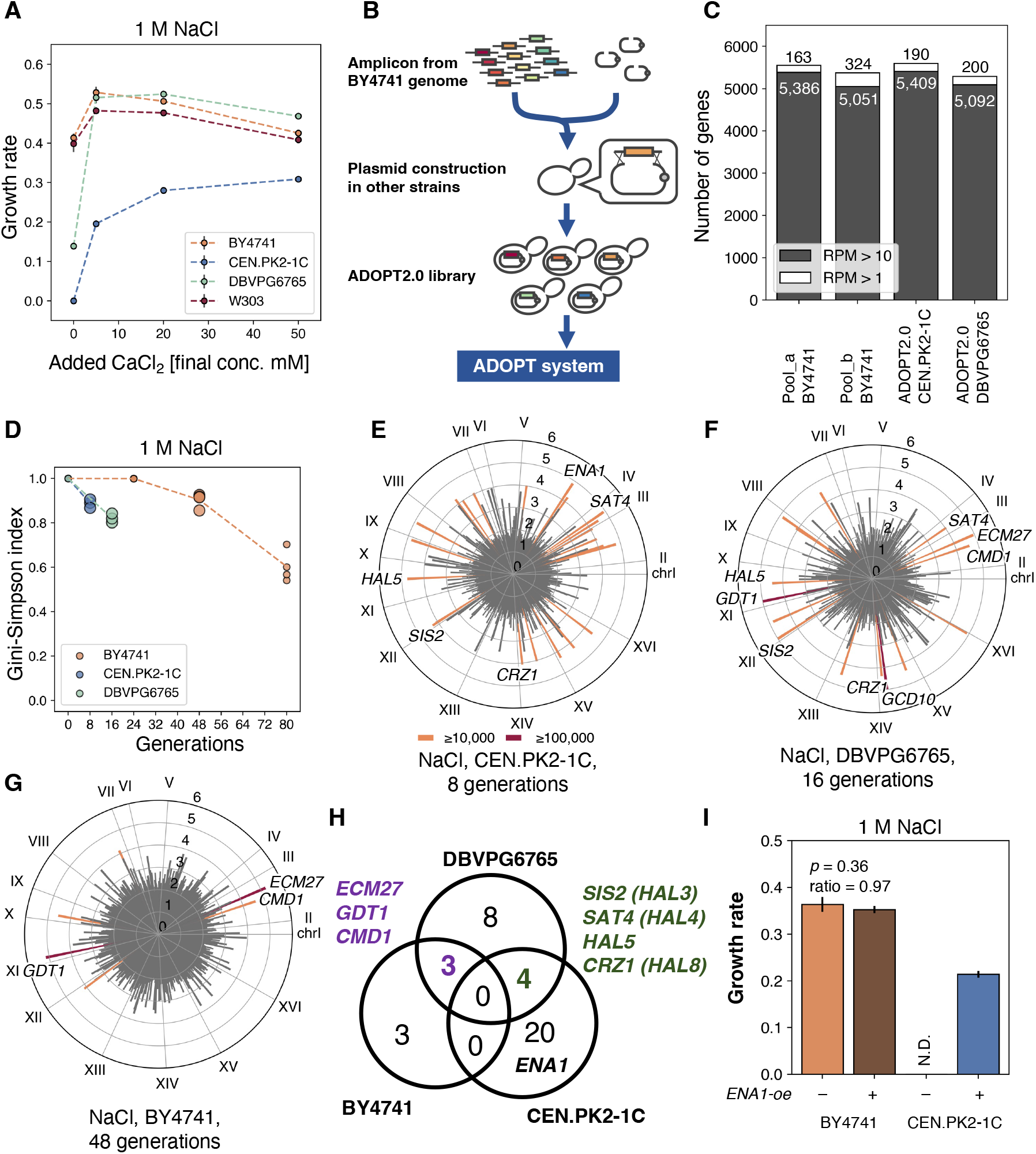
GOFAs reflect differences in yeast strains. **A)** Relationship between CaCl_2_ addition and growth rates of various *S. cerevisiae* strains. Growth rates of BY4741 (orange), CEN.PK2-1C (blue), DBVPG6765 (green), and W303 (red), under 1M NaCl with added CaCl_2_ are shown. The growth rate of CEN.PK2-1C without CaCl_2_ addition could not be defined but set to 0 for convenience. **B)** The ADOPT 2.0 libraries constructed in this study. The detail is explained in the text. **C)** Coverages of ADOPT 2.0 libraries. The filled bars indicate RPM > 10, and unfilled bars indicate RPM > 1. **D)** Decrease in the diversities of plasmids in the pooled libraries of CEN.PK2-1 (blue), DBVPG6756 (green), and BY4741 (orange), during the cultivation under salt stress. The diversity was evaluated as the Gini-Simpson index. The large circles indicate the data points used in H. **E-G)** DoE of plasmid inserts after the cultivation of ADOPT 2.0 libraries under 1 M NaCl: CEN.PK2-1C with 8 generations (E), DBVPG6765 with 16 generations (F), and BY4741 with 48 generations (G). Each cycle chart shows the average DoE in three (CEN.PK2-1C and DBVPG6765) or four replicates (BY4741). Gene names of representative GOFAs are shown with red (DoE ≥ 100,000) and orange (DoE ≥ 10,000) bars, respectively. These data are summarized in Table S4. H) A Venn diagram showing overlaps of GOFAs in BY4741, DBVPG6765, and CEN.PK2-1C. Purpled genes are common to BY4741 and DBVPG6765, and greened genes are common to CEN.PK2-1C and DBVPG6765. **I)** Growth rates of *ENA1-oe* in BY4741(oranges) and CEN.PK2-1C (blues) under salt stress. The *p*-values are from Welch’s t-test (n = 3). Error bars represent SD (n = 3).

We then developed a scheme to transfer this plasmid library to other strains. We performed homologous recombination of a mixed PCR product containing the 5,803 genes of the BY4741 genome and 2µ plasmids in yeast cells (**Fig. 3B**). The constructed pooled libraries of CEN.PK and DBVPG6765 (designated ADOPT2.0) covered more than 5,000 genes (**Fig. 3B and 3C**). Both CEN.PK and DBVPG6765 pooled libraries grew faster and adapted more rapidly to salt stress than the vector control (**Extended Data Fig. 6A**). Compared to BY4741, the diversity of the pool obtained by the Gini-Simpson index decreased more quickly (**Fig. 3D**). As a result, different GOFAs were identified in each strain (**Fig. 3E-G**). Interestingly, there were no overlapping GOFAs between BY4741 and CEN.PK (**Fig. 3H, and Extended Data Fig. 6B**). Particularly, the *ENA1* and *HAL* genes were identified as GOFAs in CEN.PK (and DBVPG6765), and *ENA1-oe* in CEN.PK restored growth under 1 M NaCl (**Fig. 3I**). Thus, the strain differences could explain why these genes were not identified in the initial ADOPT experiments on BY4741. On the other hand, the ADOPT2.0 experiments could potentially reveal strain-dependent requirements.

### GOFA reflects the factors that the strain requires in each environment

The salt sensitivity of CEN.PK has been explained by the weak expression of the *ENA* gene *ENA6* ^39^. That may explain why *ENA1* was isolated as a GOFA (**Fig. 3E and 3H**). We examined how GOFAs are altered when sufficient *ENA* function is provided to CEN.PK by *ENA1-oe* since enhanced *ENA* function seems to be a primary genetic requirement for salt tolerance in CEN.PK. Therefore, we constructed a pooled library of diploid CEN.PK (referred to as ADOPT2.1) by crossing the pooled library of ADOPT2.0 constructed in the *MATa* strain with the *ENA1*-overexpressing *MATα* strain (referred to as *ENA1-coe*) (see **Fig. 4A** and **Extended Data Fig. 7**). We also constructed another ADOPT2.1 library without *ENA1-oe* as a control (**Extended Data Fig. 7**).

**Figure 4.**
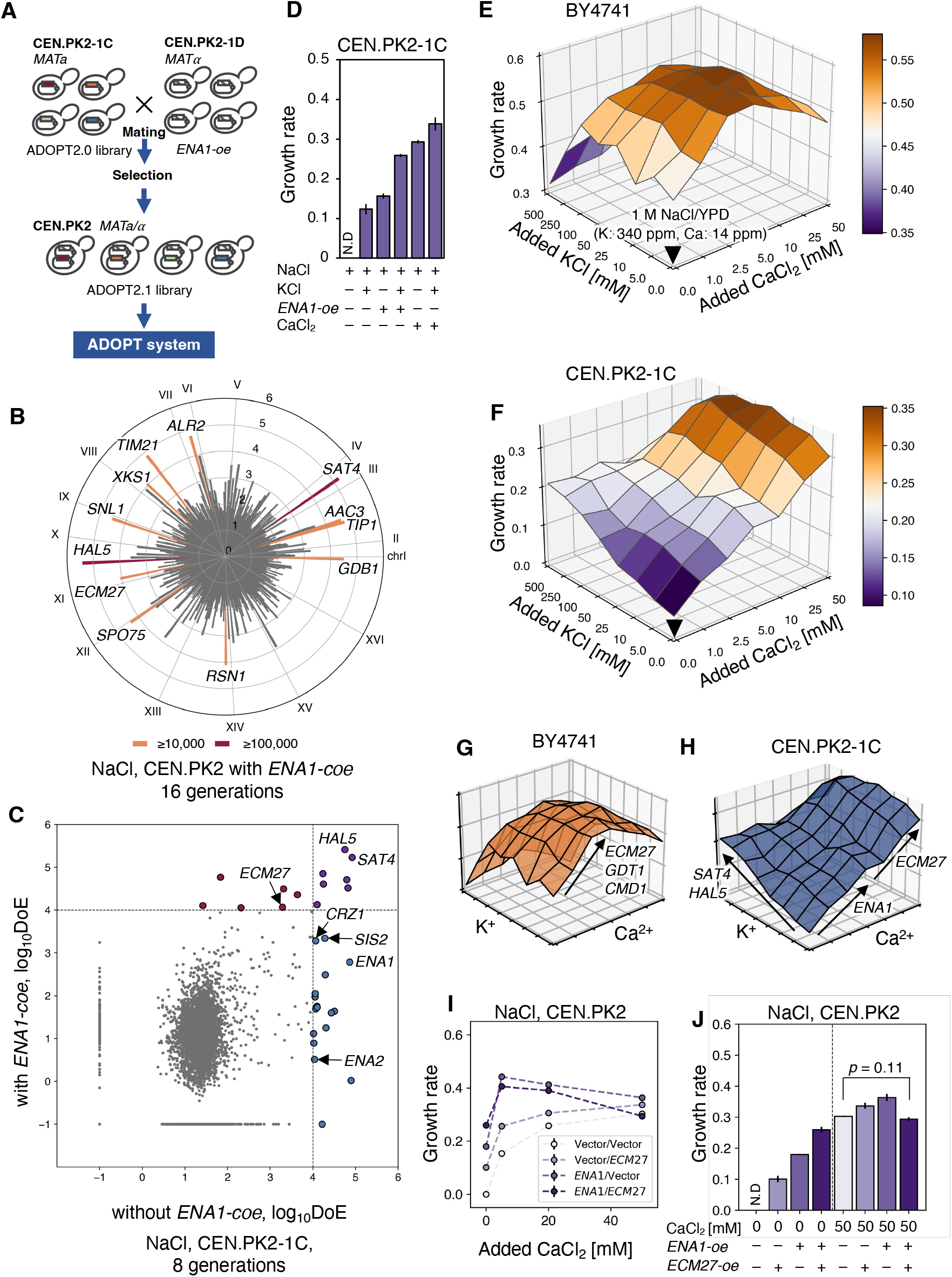
Strain-dependent requirements of calcium and potassium for the salt stress reflect strain-dependent GOFAs. **A)** Construction of the *ENA1* co-overexpression (*-coe*) ADOPT pooled library (ADOPT 2.1) by mating. The detail is explained in the text. Error bars indicate SD (n = 3). **B)** DoE of plasmid inserts after the 16 generations of ADOPT 2.1 library in CEN.PK2, under 1 M NaCl. The cycle bar chart shows the average DoE in three replicates. Gene names of GOFAs are shown with red (DoE ≥ 100,000) and orange (DoE ≥ 10,000) bars, respectively. These data are summarized in Table S4. **C)** A comparison of the average DoE with and without *ENA1-coe* under 1 M NaCl. The colored circles indicate GOFAs: without *ENA1-coe* (blue), with *ENA1-coe* (red), and both (purple). The dash lines represent the threshold of GOFAs as DoE ≥ 10,000. **D)** growth rates of CEN.PK2-1C under 1 M NaCl. N.D means not detected. Error bars indicate SD. All 15 pairs were significantly different (Welch’s t-test and Benjamini-Hochberg correction), False Discovery Rate (*FDR* < 0.05, n = 3). The value of N.D is set to 0 for the statistical test. **E-F)** Fitness landscapes of BY4741 (E) and CEN.PK2-1C (F) under 1 M NaCl with various KCl and CaCl_2_ levels. The downward triangle points to 1 M NaCl/YPD, with increasing amounts of KCl or CaCl_2_, added along the *x*- or *y*-axes. The growth rates at each KCl and CaCl_2_ addition are represented as the *z*-axis and colored as a purple-to-orange heat map, corresponding to the relative growth rate. **G-H)** A diagram of the expected relationship between slopes on fitness landscapes and GOFAs in BY4741 (G) and CEN.PK2-C (H). Arrows indicate the correspondence between Ca^2+^ or K^+^ requirement and each GOFA. **I and J)** Effects of CaCl_2_ addition on the growth rates of CEN.PK cells overexpressing *ENA1* (*ENA1-oe*) and *ECM27* (*ECM27-oe*). *ENA1 and ECM27* were overexpressed using pTOW48036 and pRS423nz, respectively. The Vector/Vector cells without CaCl_2_ addition did not grow but the growth rate was set to 0 for convenience in I and shown as N.D in J. Error bars indicate SD (n = 3). All 6 pairs with 0 mM CaCl_2_ and 5 pairs with 50 mM CaCl_2_ were significantly different (Welch’s t-test and Benjamini-Hochberg correction), *FDR* < 0.05, n = 3). A pair with no significance is shown in the figure. The value of N.D is set to 0 for the statistical test.

**Fig. 4B** shows the DoE of genes in the *ENA1-coe* pool after 16 generations under salt stress. As expected, *ENA1-oe* altered GOFAs (**Fig. 4C**). *ENA1* itself, *CRZ1*, and *SIS2* were no longer GOFA, suggesting that their functions are directly related to *ENA* function. The calcium homeostasis gene, *ECM27*, became a GOFA, suggesting calcium is a secondary requirement for satisfactory *ENA* function. The most enriched GOFAs with and without *ENA1-oe* were *SAT4* and *HAL5*, which encode protein kinases that regulate K^+^ importers ^36^. Since these GOFAs were isolated independently from the enhanced *ENA* functions, we assumed that they proposed another requirement for CEN.PK other than *ENA1*, which should be potassium. To test this possibility, we analyzed salt tolerance by adding K^+^ and found that CEN.PK could grow even under 1 M NaCl (**Fig. 4D**). The addition of K^+^ also increased the growth rate under *ENA1-oe*, and the addition of both K^+^ and Ca^2+^ further increased the growth rate (**Fig. 4D**). These results confirm that K^+^ is required for salt tolerance of CEN.PK, besides Ca^2+^ and enhanced *ENA* function.

These results strongly support the idea that GOFAs reflect factors that a given strain requires in each environment. Following this idea, the difference in salt tolerant GOFAs between BY4741 (**Fig. 1F and 3G**) and CEN.PK (**Fig. 3E and 4B**) should reflect differences in Ca^2+^ and K^+^ required for salt tolerance. Therefore, we measured the growth rates of BY4741 and CEN.PK under 1 M NaCl conditions at different Ca^2+^ and K^+^ concentrations to illuminate their respective fitness landscapes. As expected, the fitness landscapes of BY4741 and CEN.PK were markedly different (**Fig. 4E and 4F**). Ca^2+^ requirements differed between BY4741 and CEN.PK (**Fig. 4E and 4F**); BY4741 grew maximally at 5 mM Ca^2+^, while CEN.PK grew maximally at 50 mM Ca^2+^. K^+^ addition greatly increased the growth rate of CEN.PK, but not BY4741. These requirements should be imitated by overexpression of *ECM27*, *GDT1*, and *CMD1* (imitating Ca^2+^ addition) in BY4741; *ENA1* and *ECM27* (imitating Ca^2+^ addition), *SAT4* and *HAL5* (imitating K^+^ addition) in CEN.PK (**Fig. 4G and Fig. 4H**). We finally confirmed this idea for the Ca^2+^ requirement of CEN.PK; *ENA1-oe* and *ECN27-oe* additively conferred salt tolerance, but the effect was less pronounced when Ca^2+^ was added (**Fig 4I and 4J**).

Note that the Ca^2+^ and K^+^ landscapes for salt tolerance may reflect the natural conditions of the yeast. That is, salts obtained from nature contain potassium and calcium (**Extended Data Fig. 5F**), and when such salts were used, yeast growth was better than when pure NaCl was used (**Extended Data Fig. 5G**). In other words, landscapes such as those in **Figs. 4E and 4F** are likely to be optimized for the salt composition of the living environment of each strain.

### Mitochondria appear to be the primary target for enhanced salt tolerance with calcium addition

Salt tolerance in *S. cerevisiae* has been explained as the induction of *ENA* genes by the calcium pulse ^41^ (**Extended Data Fig. 8A**). However, in BY4741, Ca^2+^ addition did not enhance Ena1 protein level under 1 M NaCl (**Fig. 5A**); and *ENA1-oe* did not improve salt tolerance (**Fig. 3I**). Furthermore, Ca^2+^ addition restored growth retardation even after 15 hours of salt stress exposure (**Extended Data Fig. 8A-C**). In contrast, prior Ca^2+^ addition did not improve growth (**Extended Data Fig. 8D**). Even deletion mutants of *CNB1* and *CRZ1*, which are involved in the Ca-dependent *ENA1* induction pathway, recovered their growth rate upon Ca^2+^ addition (**Extended Data Fig. 8A and 8F**). Therefore, the induction of the *ENA* gene by short-term calcium pulse itself does not fully explain the positive effect of Ca^2+^ addition on the salt tolerance of BY4741.

**Figure 5.**
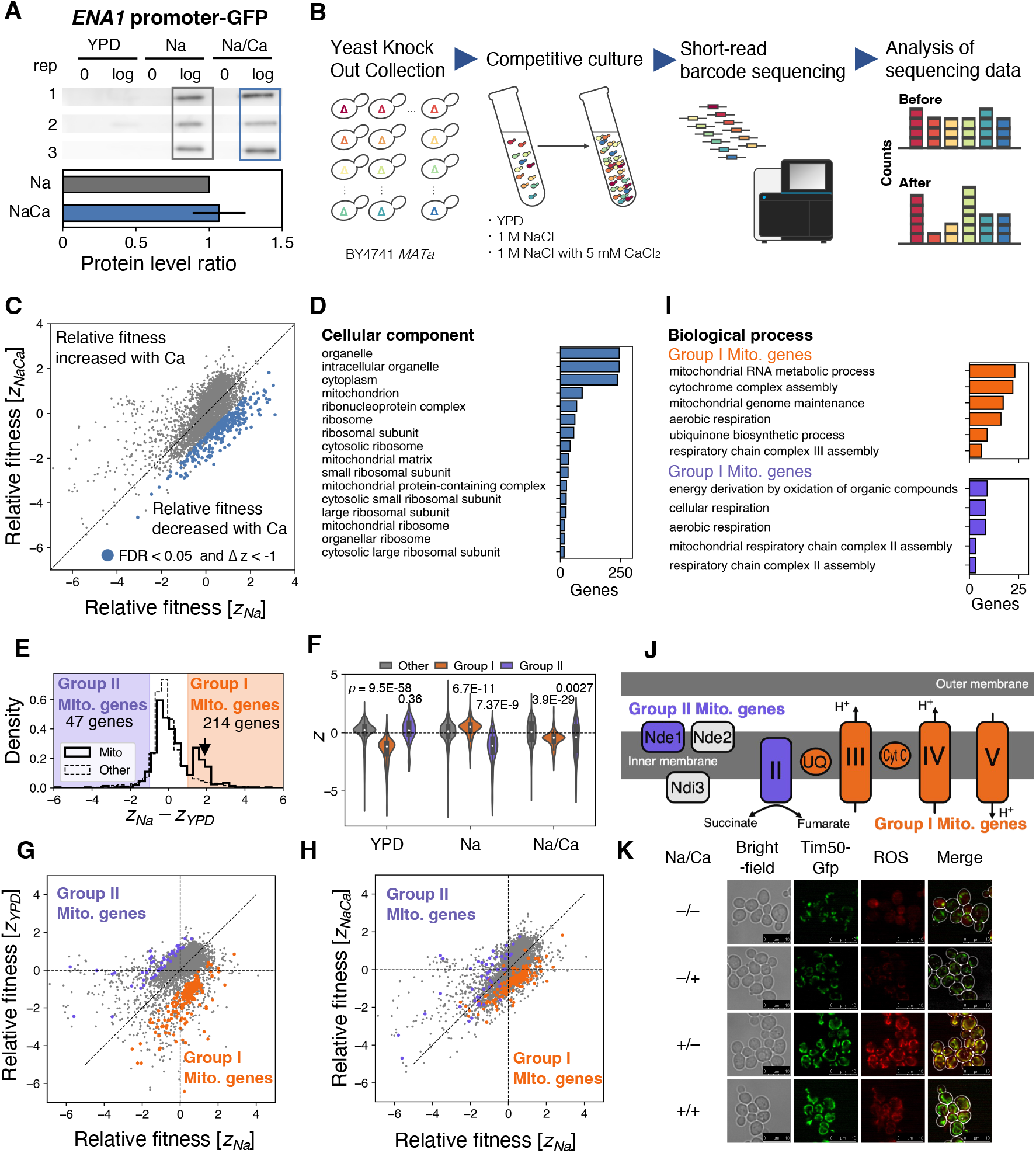
Mitochondria seem to be a prime target of enhanced salt tolerance by adding calcium. **A)** Expression of *ENA1* under the salt stress was not enhanced by CaCl_2_ addition. The *ENA1* promoter activity was detected by the Western blotting of EGFP under the control of the *ENA1* promoter under three conditions: YPD, 1 M NaCl (Na), and 1 M NaCl with 5 mM CaCl_2_/YPD (Na/Ca). The lower panel shows the EGFP level in Na/Ca relative to Na during the logarithmic growth phase. The error bar indicates the SD of relative values (n = 3). **B)** A scheme of systematic analysis for relative fitness of gene knockouts. The detail is explained in the text. **C)** Comparing relative knockouts’ fitness (Z) between Na and Na/Ca. The blue cycles indicate knockouts with reduced fitness (*FDR* < 0.05 and *ΔZ* < 1, Welch’s t-test and the Benjamini-Hochberg correction), n = 3). **D)** Enriched gene ontology (GO) terms in “cellular component” in the 296 knockouts with reduced fitness under Na/Ca (*p* < 0.05, Holm-Bonferroni correction). The bar plot shows the number of genes with indicated GO terms. Other categories of enriched GO terms are shown in Table S5. **E)** The distribution of fitness was corrected by YPD (*Z_Na_* – *Z_YPD_*). The solid and the dashed line indicate mitochondria (Mito) genes and the other genes, respectively. The orange area represents Group I Mito. genes (*Z_Na_* – Z_YPD_ ≥ 1), and the purple area means Group II Mito. genes (*Z_Na_* – *Z_YPD_* ≤ –1). **F)** The distribution of relative knockouts’ fitness of Group I (orange), Group II (purple), and the others (grey, 4,052 genes) under each condition. The *p*-values are from Welch’s t-test by comparison with Other. **G-H)** Comparisons of relative knockouts’ fitness between Na versus YPD (G) and Na versus Na/Ca (H). The purple and the orange cycles indicate the knockouts belonging to Group I and Group II, respectively. The vertical and horizontal dashed line means *Z* = 0. **I)** Enriched GO terms in “biological function” of the knockouts belonging to Group I (upper, orange) and Group II (bottom, purple) (*p* < 0.05, Holm-Bonferroni correction). A complete set of enriched GO terms are shown in Table S6. **J)** The Group I and Group II Mito. genes have separated functions in the mitochondrial respiratory chain. The diagram shows the mitochondrial respiratory chain in which complexes or proteins within Group I and Group II Mito. genes were colored orange and purple, respectively. **K)** Microscopic images of the cells with the mitochondria and their reactive oxygen species (ROS) level under four conditions. Plus, or minus of “Na” indicate YPD with or without 1 M NaCl, and plus or minus of “Ca” indicates with or without 5 mM CaCl_2_. The green color shows mitochondria inner membrane observed with Tim50-GFP. The red color indicates the mitochondrial ROS level stained by MitoTracker™ Red CM-H2Xros.

To elucidate the unknown Ca-induced salt tolerance mechanism, we used a complementary approach to ADOPT: functional profiling of gene knockout mutants. The pooled knockout collection ^42,43^ was competitively cultured and subjected to relative fitness analysis under three conditions (no salt, 1 M NaCl (Na), 1 M NaCl with 5 mM CaCl_2_ (Na/Ca)) to systematically assess gene contribution (**Fig. 5B**). The salt tolerance of knockout mutants involved in the assumed Ca-dependent mechanism should not be enhanced even by Ca^2+^ addition; in other words, the relative fitness should be lower under Na/Ca conditions than in Na conditions (indicated by the blue dots in **Fig. 5C**). We isolated 296 genes with lower relative fitness in the Na/Ca environment compared to Na (*FDR* < 0.05, *ΔZ* < –1) and found that they were enriched in the GO terms “mitochondria-related genes” and “ribosome” (**Fig. 5D, Table S5**).

We thus focused on the mitochondrial (Mito) gene. First, we examined the salt tolerance of the knockout mutants of the Mito genes and noticed their interesting behaviors; one group of Mito genes constituted a distinctly separated group with improved salt tolerance (*Z_Na_* – *Z_YPD_* >1, **Group I Mito. genes, Fig. 5E**). They appear to be salt tolerant, but this is not the case. The reasons for this can be explained as follows: (1) Under non-stress conditions, these mutants have poor relative fitness (*Z_YPD_*) due to their poorer proliferation than other mutants (**Fig. 5F and Extended Data Fig. 9A, YPD**). (2) Conversely, under salt stress, these mutants do not change their proliferation (already impaired), while most other mutants proliferate poorly. Their relative fitness (*Z_Na_*) is thus the same as the other mutations (**Fig. 5F and Extended Data Fig. 9A, YPD**). (3) The change in relative fitness between *Z_Na_* and *Z_YPD_* makes these mutants appear salt tolerant (**Fig 5E**). Ca^2+^ addition reduced their relative fitness (*Z_NaCa_*) because they did not respond to Ca^2+^, whereas the other mutants did (**Fig. 5F and Extended Data Fig. 9A, Na/Ca, and 5H**). In summary, the knockout mutants of the Groupe I Mito genes grow slowly under non-stress conditions, but their growth is unchanged under salt stress, and they do not respond to Ca^2+^ addition. Therefore, their normal function should be attacked by salt stress and restored by Ca^2+^ addition. We also noticed that some mito genes (*Z_Na_* – *Z_YPD_* < –1, **Group II Mito. genes**) showed the opposite behavior to the Group I mito genes (**Fig. 5E-H**). Group II’s growth is normal under non-stress conditions, but their growth is strongly attenuated under salt stress, and they normally respond to Ca^2+^ addition (**Fig. 5F and Extended Data Fig. 9A**). Thus, their normal function should protect the cells from salt stress, but when Ca^2+^ is sufficient, their protective function is not necessary.

We next characterized the Group I and II Mito genes in detail. Interestingly, the molecular functions of the proteins encoded in Group I and II were clearly separated between complex II and ubiquinone (UQ) of the mitochondrial respiratory chain (**Fig. 5I and J**). From this, we speculated that salt stress causes respiratory chain dysfunction (perhaps a process later than UQ), which is restored by Ca^2+^ addition. We thus observed mitochondria under salt stress with and without Ca^2+^ (**Fig. 5K**). As a result, mitochondria were more developed and generated more reactive oxygen species (ROS) under salt stress, while the addition of Ca^2+^ maintained mitochondrial development but suppressed ROS generation. Taken together, we concluded that our assumed Ca^2+^-dependent salt tolerance mechanism is related to mitochondrial function. Because of the high energy demand under salt stress, yeast cells probably need more productive mitochondrial function. As a result, high concentrations of ROS are generated, causing growth defects, which may be suppressed by the addition of Ca^2+^.

To confirm the adverse effects of salt stress on mitochondria, we investigated salt stress-induced transcriptome changes (RNA-seq) in the vector control and *CMD1-oe, ECM27-oe*, and *GDT1-oe*. The gene groups whose expression levels were significantly upregulated under salt stress conditions included “response oxidative stress (GO:0006979)”, and “arginine biosynthetic process (GO:0006526)” (**Extended Data Fig. 9B**). Under salt stress conditions, arginine uptake was significantly reduced, only about 20% (**Extended Data Fig. 9C**). Since arginine synthesis is closely related to mitochondrial function ^44^ (**Extended Data Fig. 9D**), we speculated that the elevated expression of these genes might be due to mitochondrial dysfunction under salt stress, which results in a reduction in arginine synthesis. In fact, the expression of these groups of arginine synthesis genes was commonly suppressed by *CMD1-oe*, *ECM27-oe*, and *GDT1-oe* (**Extended Data Fig. 9E**). These results are consistent with our idea that mitochondrial function is impaired under salt stress but is restored when calcium is supplied.

### Enhanced mitochondrial function can confer salt tolerance only when sufficient calcium is supplied

If the enhanced mitochondrial function is necessary for salt tolerance in yeast, why have no mitochondria-related genes been identified as GOFAs? Perhaps sufficient Ca^2+^ is required to suppress “mitochondrial runaway” under salt stress, as shown in **Fig. 5K**. If this idea is correct, mitochondria-associated GOFAs should be isolated under Ca^2+^-induced salt stress, which was the case (see below). We performed ADOPT experiments under 1 M NaCl with 5 mM CaCl_2_ (**Fig. 6A**). Upon Ca^2+^ addition, *CMD1*, *GDT1*, and *ECM27* did not become GOFAs, further supporting the idea that these GOFAs complement the Ca^2+^ requirement. On the other hand, a completely different group of genes (*CTR1*, *HAP4*, and *USV1*) were enriched as GOFAs (**Fig. 6B and 6C**). *HAP4* and *USV1* have been reported as transcription factors for mitochondrial respiratory genes ^45,46^. Of the 30 genes with genetic interactions with *HAP4* ^44^, 27 (PCC > 0.2) belong to Group I Mito genes (**Fig. 6E**), strongly suggesting a functional relationship between mitochondrial respiration and *HAP4*.

**Figure 6.**
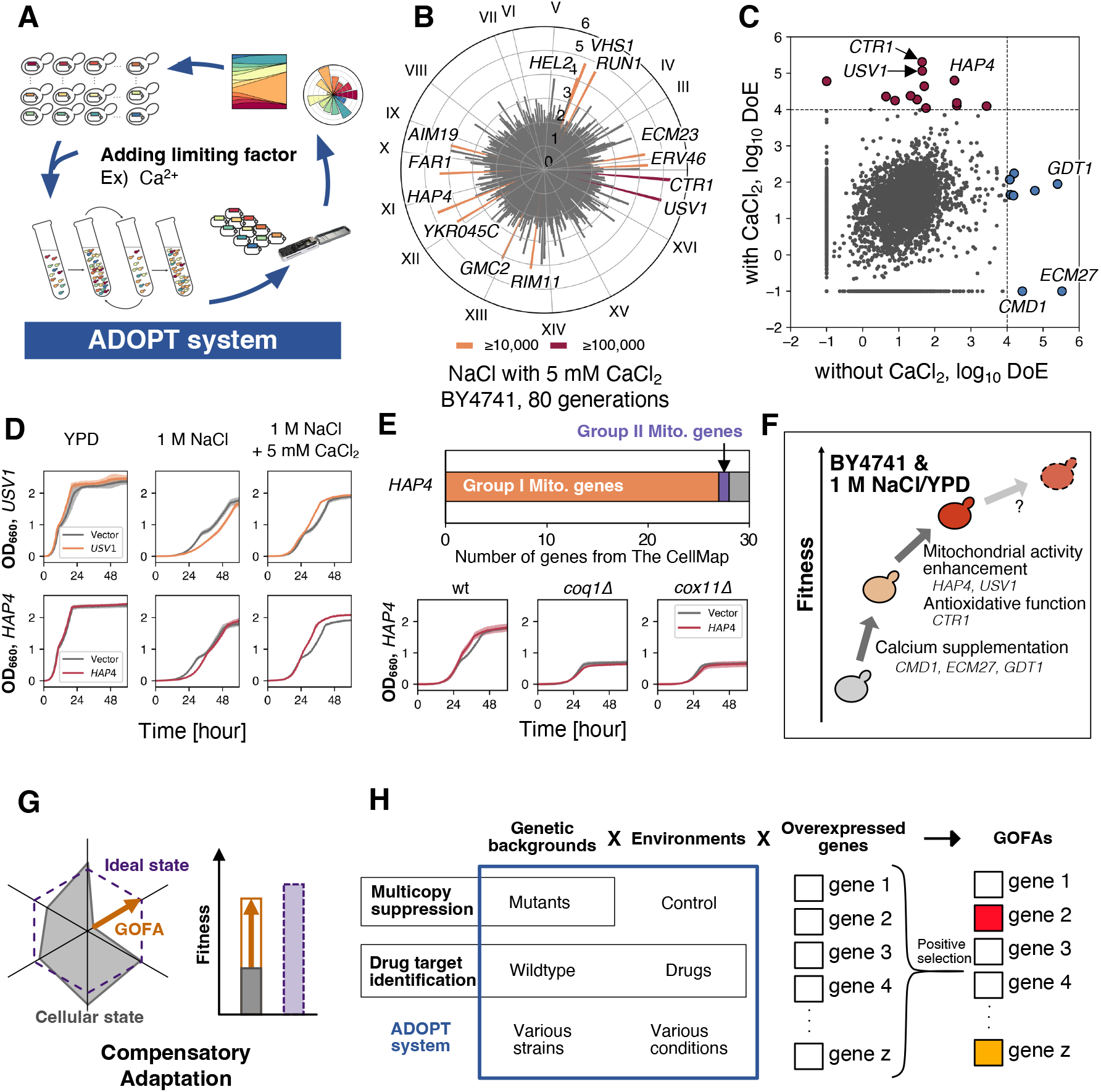
Enhancing mitochondrial function can confer salt tolerance only when enough calcium is supplied. **A)** Repeated isolation of GOFAs after the addition of a limiting factor CaCl_2_ under the salt stress. **B)** DoE of plasmid inserts after the 80 generations-cultivation of ADOPT 1.0 library (BY4741) under 1 M NaCl with added 5 mM CaCl_2_. The cycle bar chart bars show the average DoE in four replicates. The gene names of GOFAs are shown with red (DoE ≥ 100,000) and orange (DoE ≥ 10,000) bars, respectively. **C)** A comparison of the average DoE with and without CaCl_2_ addition. The colored circles indicate GOFAs, with (red) and without CaCl_2_ addition (blue). The dash lines mean the threshold of GOFAs as DoE ≥ 10,000. **D)** Growth curves of *USV1-oe* (upper, orange) and *HAP4-oe* (lower, red) under three conditions: YPD, 1 M NaCl, 1 M NaCl with 5 mM CaCl_2_. The grey line shows empty vector control. The filled areas indicate standard deviation (n = 3). **E)** The upper panel: Most genes harboring genetic interaction with *HAP4* belong to the Group I Mito. genes (PCC > 0.2, data obtained from The CellMap ^44^. The lower panel: Growth curves of *HAP4-oe* in the deletion mutant of *COQ1* and *COX11*, the Group I Mito. genes harboring the genetic interaction with *HAP4*. The filled areas indicate SD (n = 3). **F)** An illustration showing the mechanism of fitness increase of BY4741 under 1 M NaCl/YPD by fulfilling its requirements by GOFAs. **G)** GOFAs manifest the potential for cellular stress tolerance (ideal state) due to compensating for cellular requirements. **H)** Overexpression profiling using ADOPT system. The detail is explained in the text.

We confirmed that *USV1-oe* and *HAP4-oe* promoted growth under salt stress only when Ca^2+^ was supplied (**Fig. 6D**). Interestingly, overexpression of these genes without Ca^2+^ delayed growth under salt stress (**Fig. 6D**), supporting the idea that enhanced mitochondrial function under salt stress without Ca^2+^ is detrimental. Knockout of Group I Mito. genes such as *COQ1* and *COX11* did not show a *HAP4-oe* advantage (**Fig. 6E**), possibly because of *HAP4* functions upstream of these genes. These results suggest that enhanced mitochondrial activity can confer salt tolerance only when sufficient calcium is supplied (**Fig. 6F**).

Finally, we focused on *CTR1* (**Fig. 6C**). *CTR1*, encoding a copper importer ^46,47^, was isolated as the major GOFA under oxidative stress (**Fig. 1G**). *CTR1-oe* or adding 1 mM CuSO_4_ suppressed growth defects under oxidative stress (**Extended Data Fig. 10A and B**). Furthermore, instead of *CTR1*, the catalase genes *CTT1* and *CTA1* became GOFAs under oxidative stress supplied with 1 mM CuSO_4_ (**Extended Data Fig. 10B-D**). These results suggest that copper is a major limiting factor for oxidative stress and that even supplying Ca^2+^ under salt stress requires sufficient antioxidative function (**Fig. 6F**).

## Discussion

In this study, we aimed to understand the contribution of overexpression to overcoming stressful environments. Therefore, we developed a new experimental system, ADOPT, to systematically isolate genes whose overexpression is functionally adaptive (GOFAs). We first examined the characteristics of genes that become GOFAs under environmental stress. The results revealed that GOFAs are genes that compensate for cellular deficiencies and that their adaptive function strongly depends on genetic background and environment. For example, GOFAs isolated under salt stress were associated with calcium homeostasis, and their adaptive function emerged from the lack of Ca^2+^ in the medium (**Fig. 1 and 2**). In fact, under Ca^2+^-supplemented salt stress conditions, those adaptive functions were lost (**Fig. 2**), and GOFAs different from those without Ca^2+^ supplementation were isolated (**Fig. 6**). In CEN.PK, genes such as Na+ exporter *ENA1* and regulators of K^+^ homeostasis, *SAT4,* and *HAL5*, were identified as GOFAs under salt stress, but not in BY4741 (**Fig. 3**). This difference in GOFAs can be explained by the difference in Ca^2+^ and K^+^ requirements between BY4741 and CEN.PK. In fact, the adaptive effects of *ENA1-oe* and K^+^ were more substantial in CEN.PK than in BY4741 (**Fig. 4**). Based on these facts, we propose that GOFAs compensate for the missing elements for cells to reach maximum stress tolerance (”ideal state” in **Fig. 6G**). In other words, examining GOFAs reveals the missing elements necessary to maximize cellular fitness within a given genetic background and environment.

We believe that in the isolation of GOFAs, we observe a “compensatory adaptation/evolution” of cells to deficiency ^46,47^. Besides screening for drug targets, genes that fall under GOFAs in this study are often explored as multicopy repressor genes, i.e., genes that suppress/compensate the deleterious phenotype of a mutant by multicopy plasmids ^48,49^. In addition, in deleterious mutants subject to intense selection pressure, aneuploidy and consequent overexpression often occur to suppress/compensate for the harmful effects ^46,50^. On the other hand, the ADOPT experiment provided a new means of observing the complementary adaptation/evolution of various strains against potential defects under multiple environments (**Fig. 6H**).

We believe that GOFAs compensate for cellular deficiencies to achieve a potential cellular stress response. In other words, GOFAs merely augment the existing stress response system. This approach does not explain why cells have evolved sophisticated stress responses or how they acquire new stress responses. This study focused only on genes that could explain the adaptive mechanisms. In fact, among GOFAs, there are also genes whose adaptive mechanisms cannot be easily described and “emerging genes” ^27,51^ such as *YBR196C-A* in **Fig.2**. The mystery of the evolution of novel stress response mechanisms may be hidden in these genes. It may also be that the byproducts of compensatory adaptation to the environment appear as functional novelties, like the morphological novelties that occur in the compensatory evolution of gene loss ^52^.

Through the identification of GOFAs under salt stress by ADOPT, we found that calcium has a positive effect on long-lasting salt stress, distinct from the previously known short-time response of salt stress (**Fig.2 and Extended Data Fig. 8**). Furthermore, functional profiling of gene disruption mutants revealed that mitochondrial runaway might be subject to suppression by Ca^2+^ (**Fig. 5**). Overexpression of GOFAs (*HAP4* and *USV1*) identified under salt stress with supplied Ca^2+^ seem to enhance mitochondrial function, worked positively for salt stress tolerance only under calcium-supplying conditions, but rather negatively under calcium-limiting conditions (**Fig. 6**). This dictates that the primary function of calcium is to regulate mitochondrial activity under salt stress. As shown in this series of experiments, the advantage of ADOPT is its ability to rapidly and efficiently obtain GOFAs in various strains and environmental conditions (**Fig. 6H**). By using ADOPT, or “overexpression profiling,” we can uncover previously unexplored mechanisms of cellular adaptation.

## Materials and Method

### Strains and plasmids

The strains and plasmids used in this study are listed in **Table S1**.

### Medium and yeast transformation

Yeast culture and transformation were performed as previously described ^53^. We used two types of mediums: YPD and Synthetic Complete (SC) medium. YPD included 10 g/L Bacto Yeast extract (BD, USA), 20g/L Bacto Peptone (Gibco, USA), and 20 g/L D-glucose. SC medium included 6.7g/L Yeast Nitrogen Base with Ammonium Sulfate (MP, USA), 0.65 g/L DO supplement-HisLeuUra (Clontech, USA), and 20g/L D-glucose or, where appropriate, 20mg/L Histidine, 8 mg/L Uracil, and 100 mg/L Leucine. D-glucose solution was added to the medium after autoclave. Milli-Q water (Merck, Germany) was used to condition the medium. In Fig. 6D and 6E, YPD and 1 M NaCl/YPD were diluted four times with sterile water or 1 M NaCl solution. We used Shio (Shiojigyo, Japan) and Setonohonjio (Ajinomoto, Japan) as the table salt and the crude salt representatively.

### Plasmid and strain construction

RNAseq was performed as described previously ^54^. The plasmids and strains were constructed by homologous recombination activity in yeast cells ^55^, and their plasmid construction was verified by DNA sequencing.

### Exploring well-studied stress

Using the API on PubMed provided by NIH, efetch of E-utilities, we obtained 308,970 and 20,460 articles (as of 20th May 2022) searched for “yeast” and “yeast stress” (including authors. title, abstract, year of publication, and journal title), respectively. Using the TF-IDF method, we extracted the keywords with the highest scores up to 5th place from the abstract obtained for “yeast stress”^56^. Among all keywords, keywords including “stress” were extracted. Next, we determined whether or not each stress keyword appeared in the abstract of the articles obtained by “yeast stress. We used NLTK (3.6.7) for these analyses, a python library for natural language processing ^57^.

### Record growth curves and calculate growth rates

Target strains were inoculated into 4 ml of SC (–Ura or -HisUra) medium in test tubes and incubated overnight at 30°C as pre-cultivation. Then, 25 µl of pre-cultured medium were inoculated into 6 ml of target medium in L-shaped tubes and cultured at 30°C (excluding heat stress), recording optical density (OD) at 660 nm every 10 minutes with an ADVANTEC TVS062 (ADVANTEC, Japan) with shaking at 70 rpm. Growth rates [1/hour] were calculated from recorded OD data as the reciprocal of the mean doubling time, which was the slope of log base 2 of OD_660_ between 0.125 and 0.500 by linear approximation with scipy.optimize.curve_fit in the python library. If OD_660_ did not exceed 0.125 48 hours after inoculation, we designated not detected (N.D).

### Pooling gTOW6000 collection as ADOPT1.0 library

Five µl of thawed gTOW6000 collection ^17^ was inoculated into sixty 96-well plates with 200 µl of SC-Ura medium and incubated at 30°C for 48 hours. All cultured mediums were then pooled in sterile flasks and divided into 50 ml tubes. Finally, pooled libraries were with the addition of final conc—7% v/v DMSO and stored at −80 °C.

### ADOPT system

Competitive culture and passage were performed following.1 ml of an ADOPT library was inoculated into 5 ml of SC-Ura medium and incubated at 30°C overnight with shaking. Then, 24 µl (1:250 dilution) of the pre-cultured medium was inoculated into 6 ml of target medium in L-shaped tubes and cultured until stationary phase, measuring optical density with an ADVANTEC TVS062 (ADVANTEC, Japan). 24 µl (1/250 dilution) of the pre-cultured medium was passage into 6 ml of fresh medium in L-shaped tubes. The passage was repeated 1-10 times. For high stress, we used ADVANTEC TVS062 set temperatures at 37°C or 40°C in a bio shaker BR-43FL (TITEC, Japan) set at 35°C. The all-cultured medium was transferred into a 5 ml tube, centrifuged at 15,000 rpm for 1 min, its supernatant was removed, and 1 ml of 10 v/v% DMSO water was added and suspended. The suspension was transferred to 1.5 ml tubes and stored at −80°C. For the methotrexate experiment, competitive cultures were made in 500 ml Erlenmeyer flasks with 150 ml of medium, and 4.5 µl (about 1:33,000 dilution) of the culture was inoculated and passaged.

Plasmid preparation from competitive cultured yeast was performed following. 500 µl of the thawed sample was transferred to a new 1.5 ml tube, and the remaining sample was re-stored at −80°C. The sample was centrifuged at 15,000 rpm for 1 min, and its supernatant was removed. The sample was resuspended with 250 µl of Solution 1 (1 M sorbitol, 0.1 M Na_2_EDTA (pH 7.5), and 10 µg/ml RNase) and 5 µl of 10 units/µl Zymolyase–T100 (Nacalai tesque, Japan) and incubate for 30 minutes. 250 µl of Solution 2 (0.2 M NaOH and 1% w/v SDS) was added to the suspension and mixed. Then, 250 µl of Solution 3 (3 M potassium acetate and 2 M acetic acid) was added to the suspension and vortexed. The suspension was centrifuged for 10 minutes at 15,000 rpm to precipitate the insoluble material. The supernatant was added to a spin column (QIAprep Spin Miniprep Columns, Qiagen, Germany) and centrifuged at 13,000 rpm for 1 minute. After removing the column-through effluent, 750 µl of wash buffer (QIAprep Spin Miniprep Kit, Qiagen, Germany) was added and centrifuged for 1 minute. After removing the column-through effluent, the empty column was centrifuged at 13,000 rpm for 1 minute to dry the column. The column was set on a new 1.5 ml tube. The column on the tube was incubated with 50 µl of elution buffer (QIAprep Spin Miniprep Kit, Qiagen, Germany) and allowed to stand at room temperature for 2 minutes. And The column on the tube was centrifuged at 13,000 rpm for 1 minute to extract plasmids. Finally, 1 µl of the plasmid extracted solution was used to measure plasmid concentrations with a DNA staining reagent (Qubit 1X dsDNA HS Assay Kit, ThermoFisher, USA) and a Fluorometer (Qubit4, ThermoFisher, USA), the remaining solution was stored in −20°C before subsequent usage.

Long-read sequencing for plasmid inserts was performed following. Sequencing library preparation was performed according to the manufacturer’s instructions, “Four-primer PCR protocol”, using SQK-PBK-004 (Oxford Nanopore Technologies, UK). 25 ng of purified plasmid was used as each sample, and PCR reactions were performed with half of the defined protocol and the own designed primers 5’- TTTCTGTTGGTGCTGATATTGCggcgaaagggggatgtgctg-3’ and 5’- ACTTGCCTGTCGCTCTATCTTCggaaagcgggcagtgagcgc-3’. Libraries were sequenced using GridION or MinION and MinIT (Oxford Nanopore Technologies, UK) with the flow cell MinION R9.4.1. 6-12 samples per flow cell were analyzed in multiplexing. Base-calling and demultiplexing were performed using MinKNOW (Oxford Nanopore Technologies, UK) with guppy in high-throughput mode.

Analysis of sequence data and identification of GOFAs were performing following. Sequence data (fastq format) was aligned to a reference genome sequence file (R64-1-1) of budding yeast S288C using minimap2 (2.24) ^59^ to output an alignment file sam format). Next, the alignment file was reformatted and sorted using “view −Sb” and “sort” in Samtools (1.15) ^59^ to obtain a bam format file. Then, Bedtools (2.30.0) ^60^ with “bamtobed” converted the bam format file to a bed format file. The aligned reads on gTOW6000 insert locus were extracted using “bedtools intersect” with an option “-F 0.5”. The read counts on insert locus were counted by “bedtools coverage”. Subsequent analyses were performed using python (3.8.12) with NumPy (1.21.2) and pandas (1.4.1), and visualized using IGV^61^. Reads for each insert were converted to ratio (p) or reads per million (RPM). The degree of enrichment (DoE) of plasmid inserts was calculated according to the following equation,

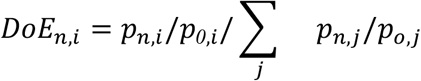

where p_0,i_ is the ratio of insert i in the pool before competitive passages and p_n,i_ is the ratio of insert i after n passages. The units of DoE are the number of plasmids per million plasmids after n passages, assuming that the initial pool was completely uniform. In this study, GOFAs were those with a DoE of 10,000 or greater. The diversity of plasmids was evaluated using the Gini-Simpson index, calculated following ^62,63^.

### Aequorin assay

Plasmid pEVP11/AEQ-HIS3 was constructed by replacing *LEU2* of pEVP11/AEQ with *HIS3* ^28^. PCR amplified the fragments of *HIS3* with primers 5’- GGCCGAGCGGTCTAAGGCGCgtttcggtgatgacggtgaa-3’ and 5’- GCGCTGGGTAAGGATGATGCgccgatttcggcctattggt-3’ using pRS413 as a template. PCR amplified the fragments of pEVP11/AEQ without *LEU2* locus with primers 5’- ttcaccgtcatcaccgaaacGCGCCTTAGACCGCTCGGCC-3’ and 5’- accaataggccgaaatcggcGCATCATCCTTACCCAGCGC-3’ using pEVP11/AEQ as a template. pEVP11/AEQ-HIS3 was introduced into each overexpressing strain. We performed transformation protocols according to Amberg 2005. Target strains were inoculated into 4 ml of SC (–Ura or -HisUra) medium in test tubes and incubated overnight at 30°C as pre-cultivation. Then, 200 µl of pre-cultured medium were inoculated into 5 ml YPD medium and cultured until OD_660_ reached 1.0. One OD_660_ unit was aliquoted into a 1.5 ml tube and centrifuged, and its supernatant was removed. The pellet was resuspended with 50 µl YPD, including 5 mM Coelenterazine H (Wako, Japan), and stood in the dark at room temperature for one hour. After centrifugation and removing the supernatant with Coelenterazine H, the pellet was washed with fresh YPD, suspended in 75 µl of YPD medium, and then applied to 96 well plates. Luminescence intensity was measured using a microplate reader MTP-880Lab (COLONA, Japan). First, fluorescence intensity was measured for 50 seconds at 5-second intervals as a baseline. Then, 25 µl of 4 M NaCl solution was added to the well by the automatic dispenser DP-50N (COLONA, Japan). The plate was agitated for 5 seconds. The fluorescence intensity was measured every 5 seconds for 10 minutes.

### Measurement of mineral concentration in the medium

We used ionometers, LAQUAtwin (Na-11, K-11, Ca-11, HORIBA, Japan), to measure the mineral concentration in the medium we used in this study. We measured following the manufacturer’s instructions. 500µl of each medium was spotted on the sensor of ionomers and measured.

### Laboratory evolutionary experiment

The culture and passages followed the “ADOPT system” described above. 1 ml of BY4741 bearing pTOWug2836 as vector control was inoculated into 5 ml SC-Ura medium and incubated at 30°C overnight with shaking. Ten passages were cultured in YPD medium containing 1 M NaCl. Genome preparation

The genome was extracted from cultured strains according to the previous report^53^ from 5 OD_660_unit cultured yeast. 500 µl of Solution 1 (1 M sorbitol Solution 1 (1 M sorbitol, 0.1 M Na_2_EDTA (pH 7.5) and 10 µg/ml RNase) and 5 µl of 10 units/µl Zymolyase solution were added to the pellet and suspended, incubated at 37°C for 30 minutes. After centrifugation and removal of the supernatant, add 250 µl buffer (20 mM Na_2_EDTA and 50 mM Tris-Cl (pH 7.4)) and 25 µl 10% SDS was added and incubated at 65°C for 30 minutes. 100 µl of 5 M potassium acetate was added to the sample and cooled on ice for 30 minutes. Then the sample was centrifuged, and its supernatant was transferred to a new 1.5 ml tube. 400 ml of isopropanol was added to the supernatant and placed at room temperature for 5 minutes. After Centrifuged again, the pellet was rinsed with 70 v/v% ethanol. 50 µl of sterile water was added to the pellet, extracting the genome. The extracted genome solution was stained with a DNA staining reagent (Qubit 1X dsDNA HS Assay Kit, ThermoFisher), and the plasmid concentration was measured with a Fluorometer (Qubit4, ThermoFisher).

Genome quality check and resequencing were outsourced to Macrogen Japan (Japan). Library preparation was performed using TrueSeq DNA PCR Free Kit (Illumina, USA), and sequencing was performed using NovaSeq 6000 (Illumina, USA) under 150 bp paired-end conditions to obtain sequence data in fastq format files. Sequence data were aligned and mapped to a reference genome sequence file (R64-1-1) of budding yeast S288C using BWA (0.7.17) ^66^. Next, the alignment file (SAM format) was converted to a bam format file and sorted using Samtools (1.15). Variants for each sample were called performed using Bamtools (1.15) with “mpileup”^65^ with “call” and filtered using vcfutils.pl varFilter (default parameters). Variants were annotated by snpEff (4.1) ^65,66^ using R64-1-1.86. Comparison of An and Ev variations was performed by bcftools isec. The raw data were available in the DNA Data Bank of Japan (accession number: DRA014470).

### Construction of ADOPT 2.0 libraries by transformation

The mixed PCR products of S288C’s ORF and the two plasmid fragments from pTOW40836 were introduced into the target strain. Transformation protocols were performed according to the previous report^53^. To 6 ml of yeast with 1 OD_660_ unit, 60 µl of PCR mix, 30 µl of plasmid fragments, 1,440 µl of 50 w/v % polyethylene glycol 4,000, 216 µl of 1 M LiOH, and 144 µl of ssDNA were added and spread on 15 cm diameter SC-Ura agar medium. Incubated at 30°C for 48 hours, scraped off the formed colonies on all agar templates, and added DMSO to final conc. 7 v/v%, and stored at −80°C. Yeast cells were equivalent to 2 agar plates for DBVPG6765 and 5 agar plates for CEN.PK2-1C was transformed.

### Construction of ADOPT 2.1 libraries by mating

The ADOPT 2.0 library in CEN.PK2-1C and CEN.PK2-1D bearing pRS423nz2-ENA1 were mixed to be 1:1. Each used 2 OD units. Mixed strains were spread and cultured on an SC-HisUra agar plate (15 cm) for 48 hours at 30°C. After incubation, to selectively reduce unmated yeasts, the colonies were scraped, and 2 OD units each were spread again on 5 plates of fresh SC-HisUra agar and incubated at 30°C for 48 hours. Finally, the colonies were scraped off, and DMSO was added to reach the final conc. 7%, and stored at −80°C.

### GFP western blot analysis

GFP was detected by western blot as described previously ^68^. ENA1-GFP cells were cultivated in YPD, 1 M NaCl/YPD and 1 M NaCl/YPD with 5 mM CaCl_2_. One OD unit of the cells was harvested at the log phase (OD_660_ = 1.0). The cells were treated with 1 ml 0.2 mol/l NaOH and then 50 µl 1xNuPAGE LDS sample buffer (Invitrogen, USA) and heated at 70°C for 10 mins. Protein lysate was labeled with Ezlabel FluoroNeo (ATTO, Japan) and separated by polyacrylamide gel electrophoresis on 4–12% on NuPAGE 4%–12% Bis-Tris Gel (Invitrogen, USA). The separated proteins were transferred onto a PVDF membrane (Invitrogen, USA) using the iBlot (Invitrogen, USA). GFP was probed by the anti-GFP antibody (Roche) (1: 1,000), peroxidase-conjugated secondary antibody (Nichirei Biosciences, Japan) (1: 1,000), and SuperSignal West Femto Maximum Sensitivity Substrate (Thermo Fisher Scientific, USA). In chemiluminescence detection mode, GFP was detected and measured using the LAS-4000 image analyzer (Fujifilm, Japan). Quantification of the band intensity was carried out using ImageJ (1.53k).

### Genetic profiling using yeast gene knockout collection

96-well plates were dispensed with 200 µl of YPD, inoculated with 5 µl of thawed Yeast Knockout Out Haploid MAT-a Collection ^69^, and incubated at 30°C for 48 hours. All culture strains were mixed in sterile flasks, divided into 50 ml tubes, and added final conc. 7 % v/v DMSO, and stored at −80 °C. One ml of Pooled KO library was inoculated into 5 ml of YPD medium and incubated at 30°C overnight with shaking. Next, 6 ml of medium was dispensed into an L-shaped tube, and 24 µl (1/250 dilution) of the pre-culture was inoculated and incubated for a fixed time until steady-state while measuring optical density with an ADVANTEC TVS062. After a particular time, 6 ml of fresh medium was dispensed into another L-shaped tube, and the culture was passaged 24 µl (1/250dilution) of the culture. The passage was repeated two times. The genome of harvested cells was extracted (see Laboratory evolutionary experiment). Strain-specific DNA barcodes were amplified using multiplex primers and a common U2 primer. PCR conditions were set as follows: 5 min at 98°C for initial denaturation, 30 cycles of 30 sec at 98°C, 30 sec at 55°C, 45 sec at 72°C, and a final extension time of 10 min at 72°C. PCR products were purified from 2% agarose gels using a Geneclean III kit (Qbiogene, USA), quantified using a Kapa qPCR kit (Sigma-Aldrich, USA), and sequenced with an Illumina HiSeq 2500 machine. Sequence analysis was performed on the second passages and the pre-culture pool. Each experiment was performed in biological triplicates performed for all conditions. We denoted relative fitness in terms of *Z*-scores, which was the standard normalized distribution of fold change between RPM of barcodes before and after cultivation. The false discovery rate (*FDR*) for the *Z*-score between conditions was calculated using Welch’s t-test ^70^ and the Benjamini-Hochberg correction ^71^. GO enrichment analysis was performed using the Gene Lists function on the SGD website (www.yeastgenome.org/).

### Microscopic observation

Microscopic observation was performed as described previously ^72^. TIM50-GFP^73^ were cultured in YPD, YPD with 5 mM CaCl_2_, 1 M NaCl/YPD, and 1 M NaCl/YPD with 5 mM CaCl_2_. Cells were harvested at the log phase (OD_660_ = 1.0), and 1 μl of the suspension cell was mixed with 2 μl of YPD on a glass slide. Images were obtained and processed using the DMI6000 B microscope and Leica Application Suite X software (Leica Microsystems, Germany). The GFP fluorescence was observed using the GFP filter cube (Leica cat. # 11513899). Mitochondria were stained with 100 nM of MitoTracker Red CM-H2Xros (M7513, Thermo Fisher Scientific, USA) for 30 min and then washed with 0.5 ml of YPD. The cells were then observed using RFP filter cubes (Leica cat. # 11513894).

### RNAseq

RNAseq was performed as described previously ^74^. The four strains: *CMD1-oe*, *ECM27-oe*, *GDT1-oe*, and vector control, were pre-cultured in SC-Ura at 30°C overnight and cultured in YPD or YPD with 1M NaCl medium and harvested at the log growth phase (OD_660_ = 1.0). Purified RNA was quality-checked by BioAnalyzer (Agilent, USA) or MultiNA (Shimazu, Japan), and concentration was measured by Qubit (Thermo Fisher Scientific, USA). Purified RNA was stored at −80°C until subsequent experiments. cDNA library was prepared using the TrueSeq Stranded Total RNA kit (Illumina, USA) and half the protocol of the TrueSeq RNA library prep kit. 4 µg of the library was prepared by adding 1 µl of 142.8x diluted ERCC RNA Spike-in mix (ThermoFisher, USA) to 4 µg of total RNA. Libraries were quality checked on an Agilent 2100 BioAnalyzer (Agilent, USA), concentrations were measured on a Real-TIme PCR system (ThermoFisher, USA), and libraries were pooled. cDNA library Sequencing was performed by pair-end sequencing on an Illumina NextSeq 550 (Illumina, USA). Three biological duplications were analyzed for all strains. The sequences were checked for sequence quality by FastP ^75^ and then aligned using Hisat2 ^76^. The aligned data were formatted into bam files by Samtools ^59^. Finally, expression level variation analysis was performed by EdgeR ^59,76^. The raw data were available in the DNA Data Bank of Japan (accession number: DRA014472). GO enrichment analysis was performed using the Gene Lists function on the SGD website (www.yeastgenome.org/).

### Arginine uptake assay

BY4741 wild-type and *can1*Δ cells were cultured in YPD or YPD with 1 M NaCl medium and harvested at the log growth phase (OD_660_ = 1.0) and suspended in 4 ml of YPD or YPD with 1 M NaCl medium at a density of 2.5 × 10^8^ cells/ml, respectively. The arginine uptake reaction was initiated by the addition of 1.0 ml YPD or YPD with 1 M NaCl medium containing [U-^14^C]arginine at final radioactivity level of 0.518 kBq/ml, respectively. Immediately after addition (defined as 0 min), and after incubation for 60 min at 30°C, 0.5 ml aliquots of cell suspension were withdrawn, and filtered on cellulose acetate membrane filters (0.45 µm; ADVANTEC, Japan), and washed with cold 10 mM HEPES (pH6.4). The radioactivities of recovered cells were measured using a liquid scintillation counter. Arginine uptake was calculated by subtracting the radioactivity of 0 min from that of 60 min. For the normalization of arginine uptake, protein contents in cells mixed with YPD not containing radiolabeled arginine and collected at 0 min and 60 min were measured by Lowry method.

### Quantification and Statistical Analysis

Information on statistical analysis and biological replicates is included in figure legends. The significance level was set at 0.05.

## Acknowledgments

We thank the members of the Moriya Laboratory (Okayama University) for their helpful discussions. We thank Nobuyuki Uozumi and Patrick Masson for sharing pEVP11/AEQ, Michael Knop for the concept inspiration, Luis Alberto Vega Isuhuaylas for chemical genomics analytical support, Hiroaki Mano, Yuuki Kobayashi, Shoko Ohi, Mika Ikeda, and Wen Xin Xuan for high-throughput sequencing support. This work was partly supported by JSPS KAKENHI Grant Numbers 18H04824 20H03242, and 20H04870 (H.M.), 21J12451 (N.S.), and JP17H06411 (C.B. and Y.Y).

## Extended Data Figure

**Extended Data Figure 1.**
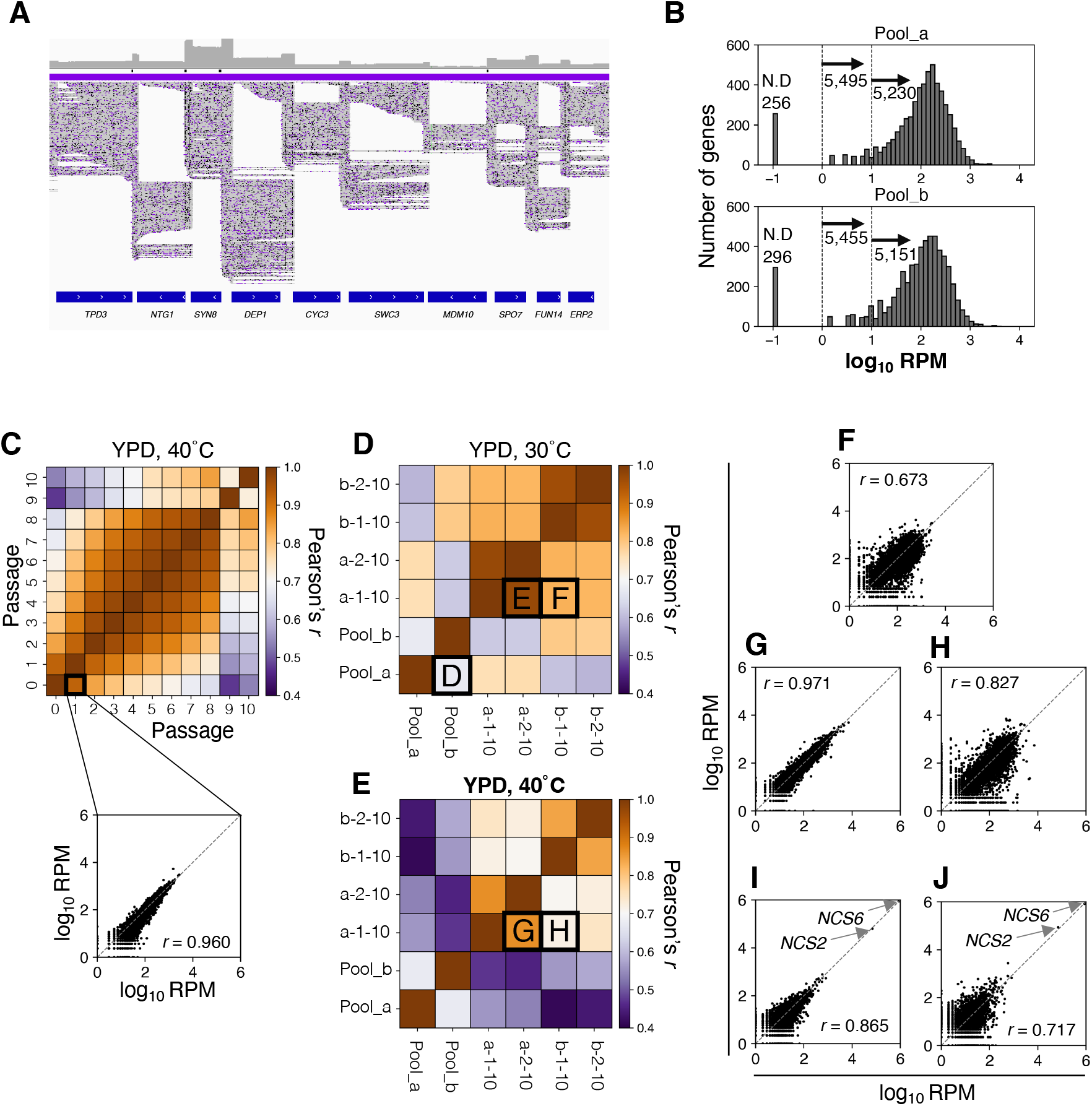
A quality check of sequencing data by ADOPT system. **A)** An example of plasmid inserts sequencing result, visualized by a genome browser, IGV. **B)** Distribution of plasmid inserts (RPM) in pool_a (upper) and pool_b (lower). Those with an RPM of 0 are appended with 0.1 for convenience. Vertical dashed lines indicate RPM >1 or RPM >10. The mean, median, and SD of RPM in pool_a were 76.8, 120, and 6.42, respectively, and in pool-b were 70.6, 115, and 7.05. **C)** A heatmap showing Pearson’s correlation among each passage in the identical replicate under 40°C. A purple-to-orange presents low to high Pearson’s correlations. The lower panel shows a scatter plot comparing RPM before and after 1^st^ passage. **D-E)** RPM scores were quite reproducible between the replicates that originated the identical pool, while they had some differences when originated pools differed. Heatmaps showing Pearson’s correlation among four replicates on the 10^th^ passages under (D) 30°C and (E) 40°C. “a-” and “b-” in the four replicates originated from pool_a and pool_b responsively. **F-J)** Scatter plots comparing RPM. The comparisons are described in D and E.

**Extended Data Figure 2.**
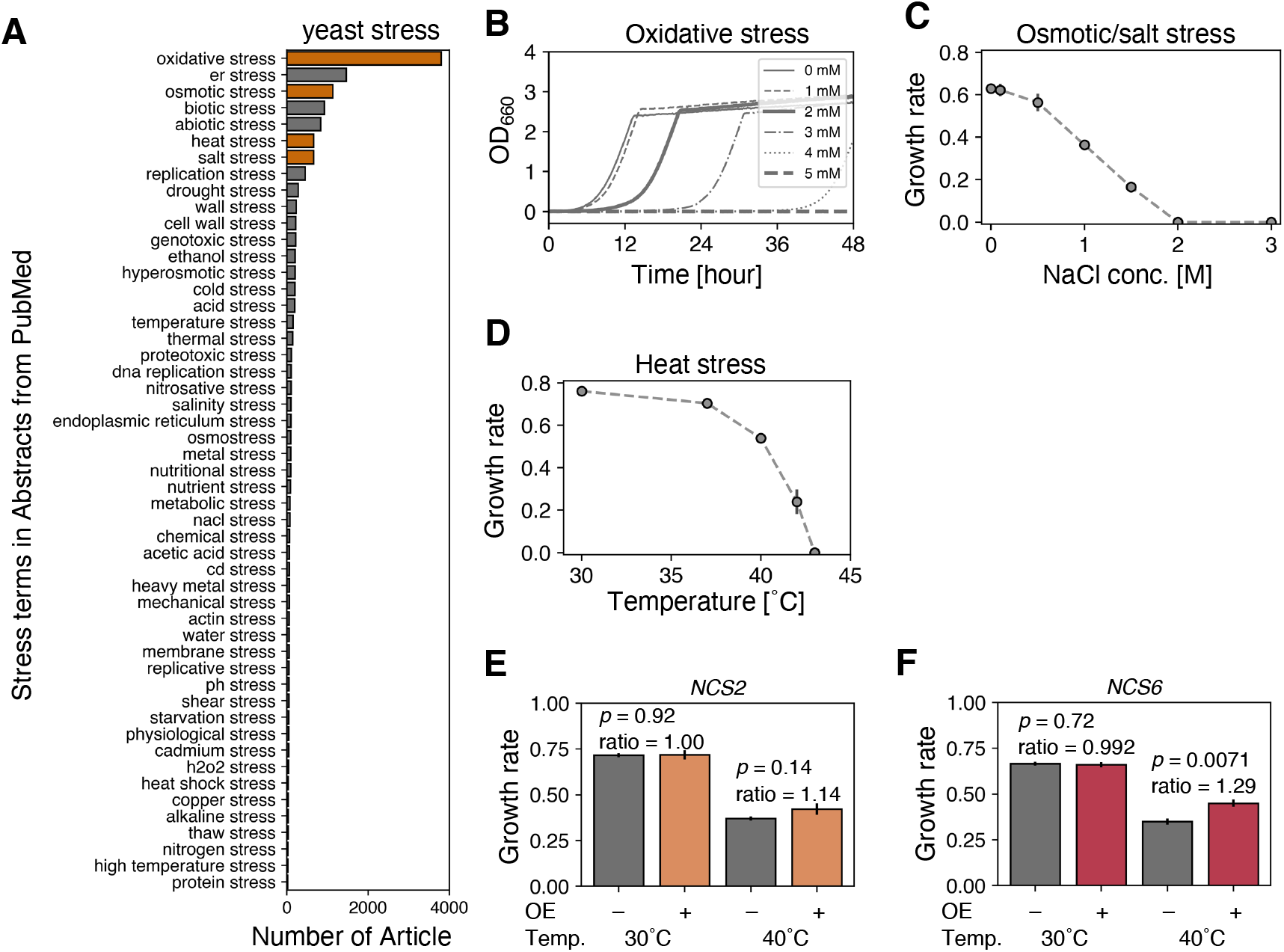
The well-studied stress in yeast. **A)** The number of articles in which “XXX stress” appears in the abstract of the articles obtained from PubMed search for “yeast stress”. The orange bars indicate the stresses which we focused on in this study. **B)** Growth curves of BY4741 with various amounts of H_2_O_2_ in YPD. **C)** Growth rates of BY4741 with various amounts of NaCl in YPD. **D)** Growth rates of BY4741 in YPD under some temperatures. **E-F)** The growth rates of BY4741 cells overexpressing (*-oe*). (E) *NCS2-oe* and (F) *NCS6-oe* at 30°C and 40°C. The *p*-values are from two-tailed Welch’s t-test (n = 3). Ratios were the average growth rate over the empty vector control. Error bars show SD (n = 3).

**Extended Data Figure 3.**
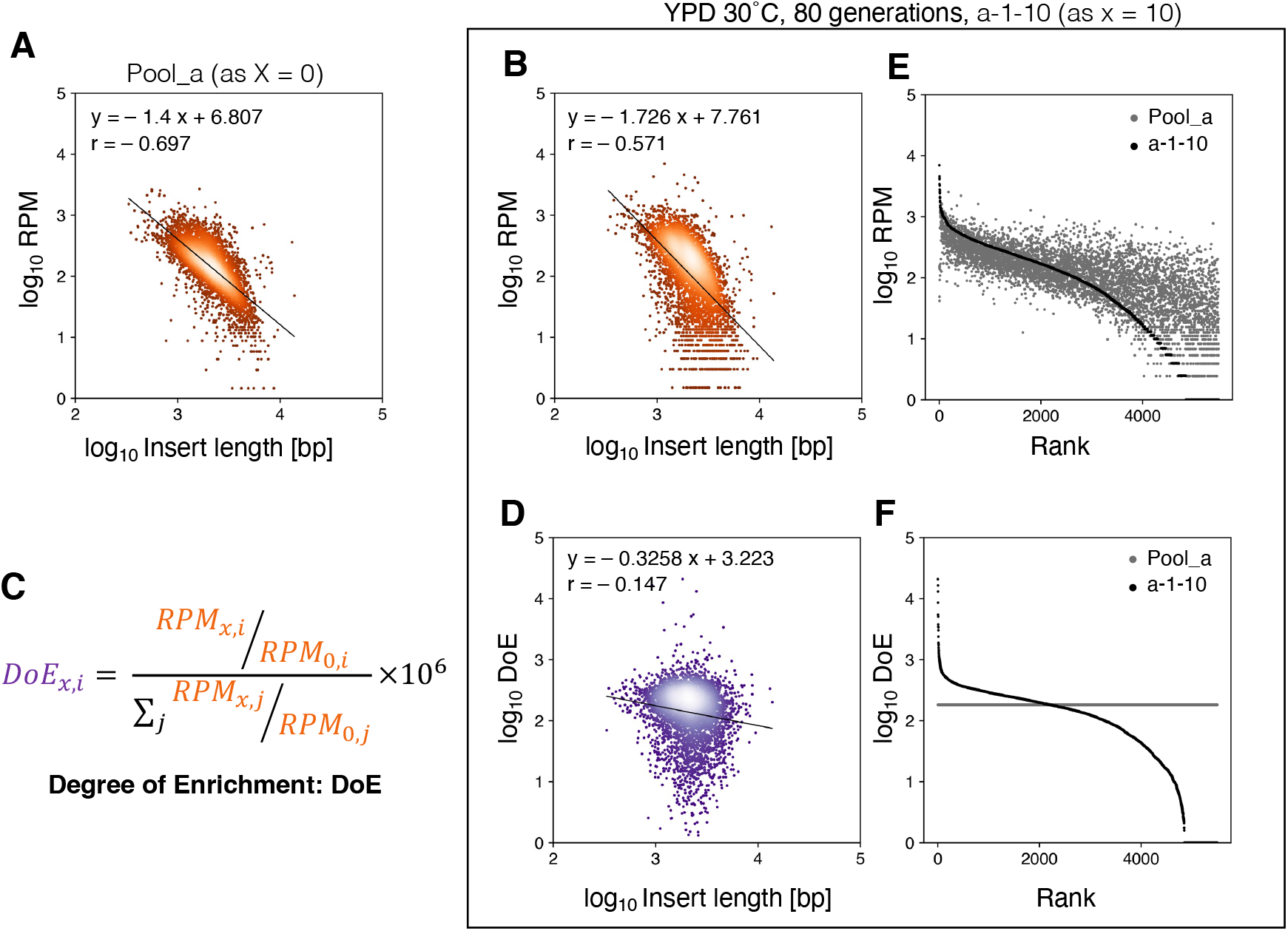
Calculating Degree of Enrichment (DoE) to be corrected for insert frequency bias by inset length. **A-B)** Negative correlations between insert frequency (RPM) and insert length (bp) in (A) pool_a or (B) 10^th^ passage. The solid lines mean regression lines (regression equations are shown in the figure.). The shading of colors implies the density of the points, which are thinner with higher density. **C)** The formula for calculating Degree of Enrichment (DoE). The orange terms mean RPM (corresponding to A as 0 and B as x), and the purple term means DoE (corresponding to D). The detail is explained in material and method. **D)** A scatter plot comparing DoE and inset length in the 10^th^ passage. The solid lines mean regression lines (regression equations are shown in the figure). The shading of colors implies the density of the points, which are thinner with higher density. **E-F)** Scatter plots show (E) RPM or (F) DoE in the initial pool (grey) and the 10^th^ passage (dark grey). These scores were sorted in descending order of RPM and DoE in the 10^th^ passage.

**Extended Data Figure 4.**
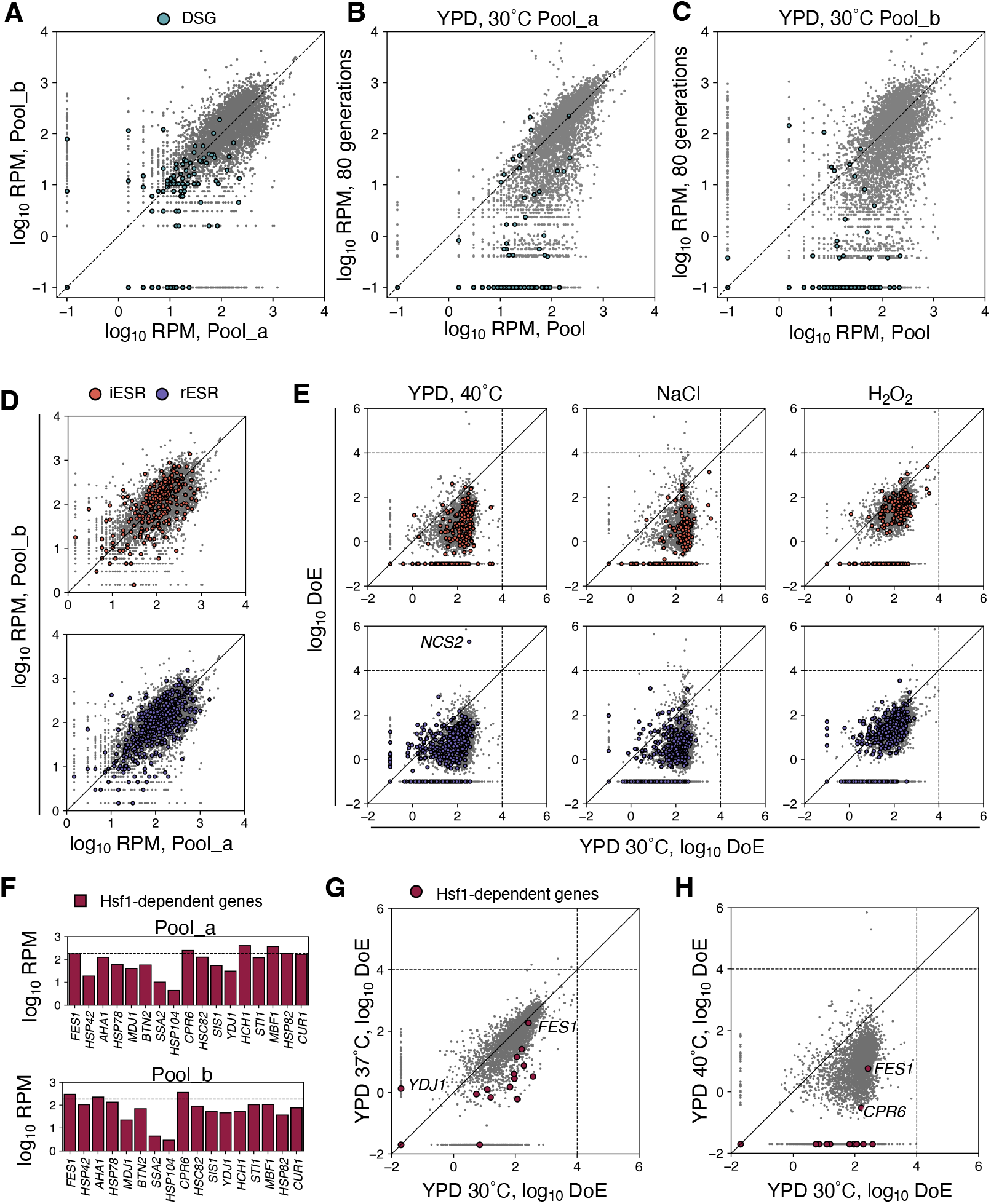
Stress-induced genes were not enriched as GOFAs. **A-C)** RPM of dosage-sensitive genes (DSG, green circle) at (A) initial in pool_a and pool_b, and (B) after 80 generation cultivation under YPD 30°C in pool_a, (C) and pool_b. **D)** Initial RPM of induced Environmental Stress Response genes (iESR, top and orange circle) and reduced ESR (rESR, bottom, and purple circle) in Pool_a and Pool_b. The grey circles mean other genes than ESR. **E)** The scatter plots show comparisons of DoE of iESR (top and orange circle) and rESR (bottom and purple circle) between 30°C and other three conditions: 40°C (left), 1 M NaCl (middle), and 2 mM H_2_O_2_ (right). The grey circles mean other genes than ESR. All circles are the mean DoE of four replicates. The horizontal and vertical dashed lines indicate the threshold of GOFAs (DoE ≥ 10,000). Genes with a DoE of 0 (undetected) have 0.1 added for convenience. **F)** Initial RPM of heat shock response genes (Hsf1-dependent genes) in Pool_a (top) and Pool_b (bottom). **G-H)** The scatter plots show comparisons of DoE of Hsf1-dependent genes between 30°C and (G) 37°C or (H) 40°C. The red and grey circles mean Hsf1-dependent genes and other genes responsively. All circles are the average DoE of four replicates. The horizontal and vertical dashed lines indicate the threshold of GOFAs (DoE ≥ 10,000). Genes with a DoE of 0 (undetected) have 0.1 added for convenience.

**Extended Data Figure 5.**
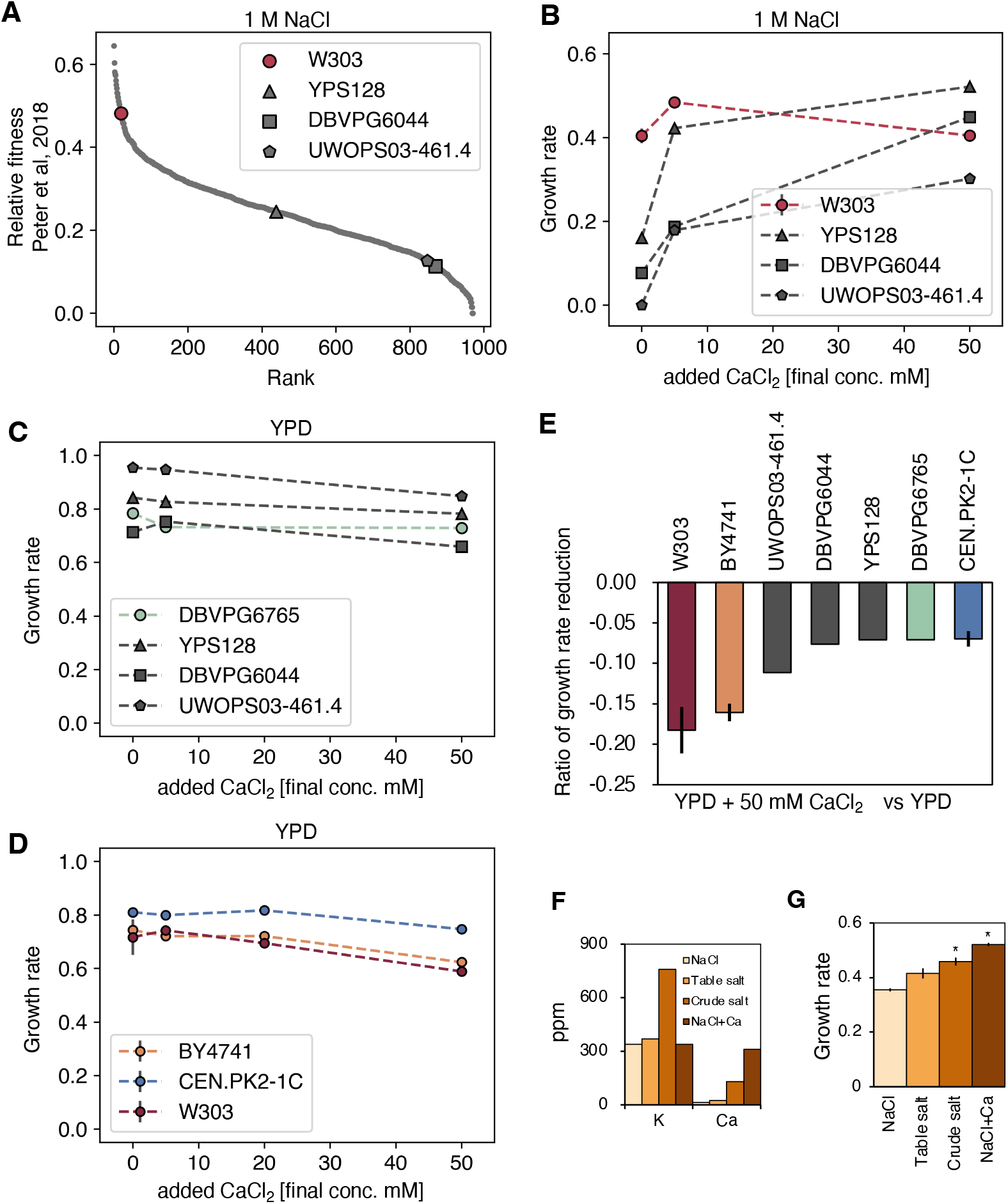
The addition of Ca^2+^ increased the growth rates of various strains under salt stress but did not without salt stress. **A)** Relative fitness of various strains; W303 (red, circle), YPD128 (triangle), DBVPG6044 (square), and UWOPS03-461.4 (pentagon), under 1M NaCl. The relative fitness data were from Peter et al. 2018. **B)** Relationship between the addition of CaCl_2_ and growth rates of various strains under 1 M NaCl. The growth rate of UWOPS03-461.4 without CaCl_2_ addition could not be defined but set to 0 for convenience. Three biological replicates were measured for W303. **C)** Relationship between the addition of CaCl_2_ and growth rates of various strains under YPD. The green circles mean DBVPG6765’s growth rates. **D)** Relationship between the addition of CaCl_2_ and growth rates of various strains; BY4741 (orange), CEN.PK2-1C (blue), and W303 (red), under YPD. **E)** The ratio of reduction in growth when 50 mM CaCl_2_ was added compared to YPD. Three biological replicates were measured for W303, BY4741, and CEN.PK2-1C. **F)** Growth rate of BY4741 under three salts: NaCl as an experiential reagent, a table salt, and a crude salt. NaCl+Ca means NaCl medium adding 5 mM CaCl_2_. Asterisks indicate significant differences compared to NaCl (Welch’s t-test and Bonferroni correction (*p* < 0.05/3)). **G)** K^+^ and Ca^2+^ concentrations in the medium used in F. Na^+^ concentrations in all mediums were adjusted to 23.0 g/l. Error bars indicate SD (n = 3).

**Extended Data Figure 6.**
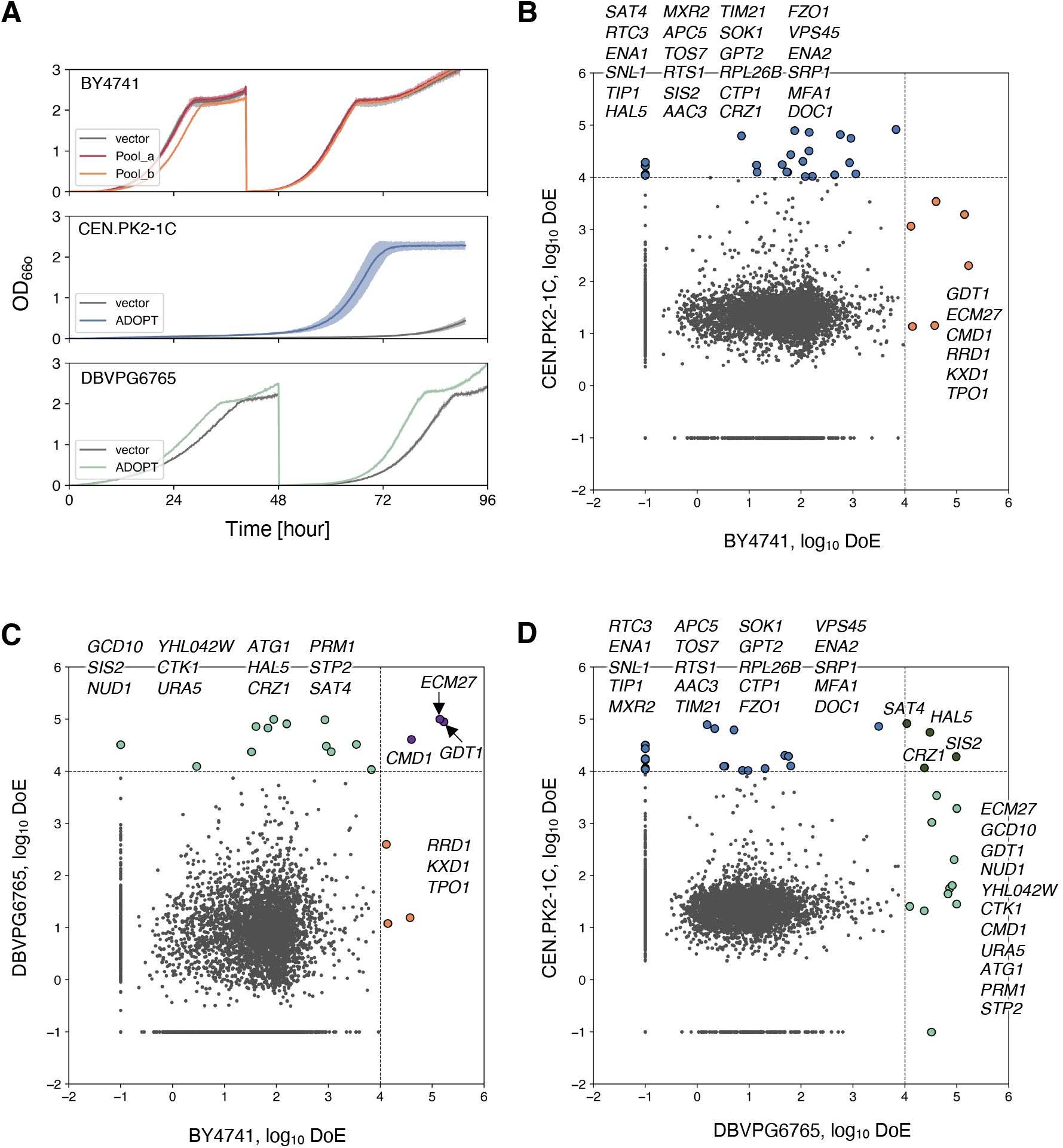
Supplement to ADOPT2.0 experiments. **A)** ADOPT 2.0 libraries of CEN.PK2-1C (middle, blue) and DBVPG6765 (bottom, light green) grew faster than the vector controls and quickly adapted to the salt stress. The solid lines and the filled areas indicate the average of OD_660_ and the standard deviation (n = 3) responsively. The grey means the empty vector controls. The top panel shows the growth curves of ADOPT1.0 libraries in BY4741 under salt stress. The red and orange correspond to replicates derived from Pool_a and Pool_b (each n = 2). **B-D)** Scatter plots show comparisons of the average DoE; between (B) BY4741 and CEN.PK2-1C, (C) BY4741 and DBVPG6765, (D) DBVPG6765 and CEN.PK2-1C, under 1M NaCl. The orange, blue, and light green circles indicate GOFAs in BY4741, CEN.PK2-1C, and DBVPG6765, responsively. The purple and green circles mean GOFAs in both BY4741 and DBVPG6765, and both CEN.PK2-1C and DBVPG6765. Genes with a DoE of 0 (undetected) have 0.1 added for convenience. The horizontal and vertical dashed lines indicate the threshold of GOFAs (DoE ≥ 10,000).

**Extended Data Figure 7.**
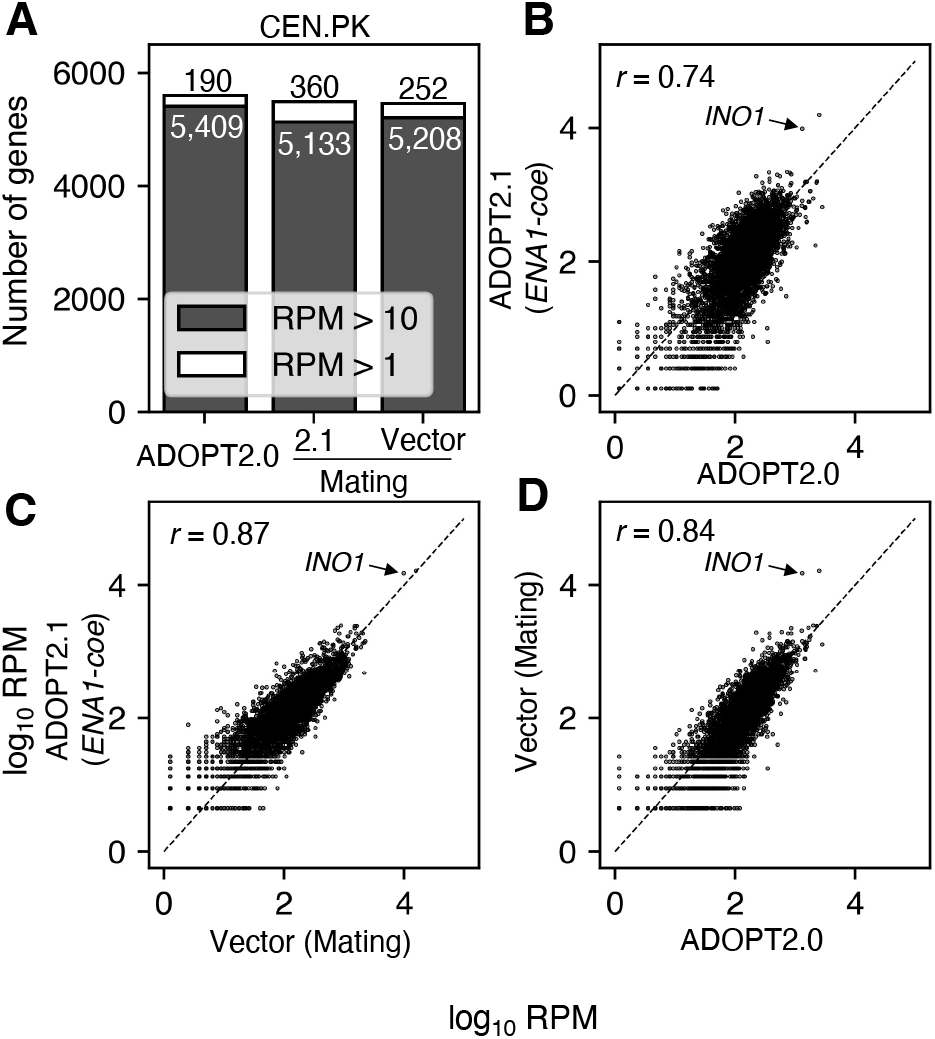
The quality check of the ADOPT2.1 library. **A)** ADOPT 2.1 libraries covered over 5,000 genes in CEN.PK. The left, middle, and right bars indicate ADOPT2.0 library, ADOPT2.1 library with *ENA1-coe,* and with empty vector as a control for mating, responsively. The filled and opened bars indicate RPM > 10 and RPM > 1. **B-C)** Scatter plots show comparisons of initial RPM between (B) ADOPT2.0 and ADOPT2.1, (C) ADOPT2.1 and Vector, and (D) ADOPT2.0 and Vector. As an addendum, we found that *INO1* was enriched in these libraries during construction and might compensate for the lack of Inositol in the SC medium used in the selection ^77^.

**Extended Data Figure 8.**
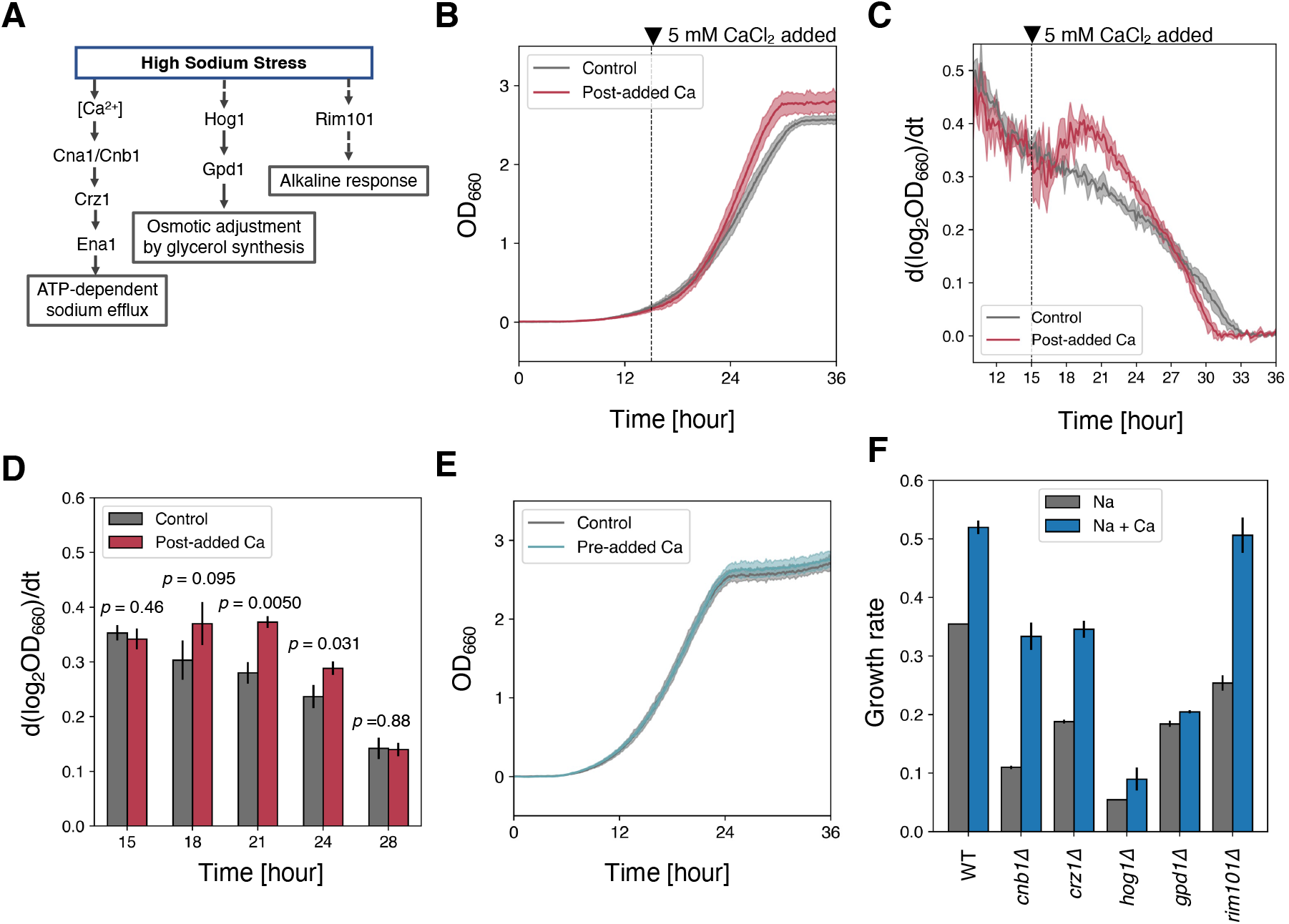
The effects of calcium addition alone cannot be explained by short-term stress response enhancement. **A)** A scheme of major responses to salt stress. B-D) Growth of BY4741 under salt stress with CaCl_2_ added after salt stress exposure. **B-D)** The growth curves of BY4741 under 1 M NaCl (grey) and post-added 5 mM CaCl_2_ at 15 hours later (red). (C) Instantaneous growth rates per hour and (D) their comparison at every 3 hours. The vertical dashed line means the timing of CaCl_2_ addition. The filled areas and error bars indicate SD (n = 3). The *p*-values are from Welch’s t-test. **E)** The growth curves of BY4741 under 1 M NaCl (grey) and pre-added 5 mM CaCl_2_ at pre-cultivation (green). The filled areas indicate SD (n = 3). **F)** Growth rates of knockouts related to major responses to salt stress. The grey and blue bars indicate growth rates under 1 M NaCl with/without 5 mM CaCal_2_ responsively. Error bars indicate SD (n = 2).

**Extended Data Figure 9.**
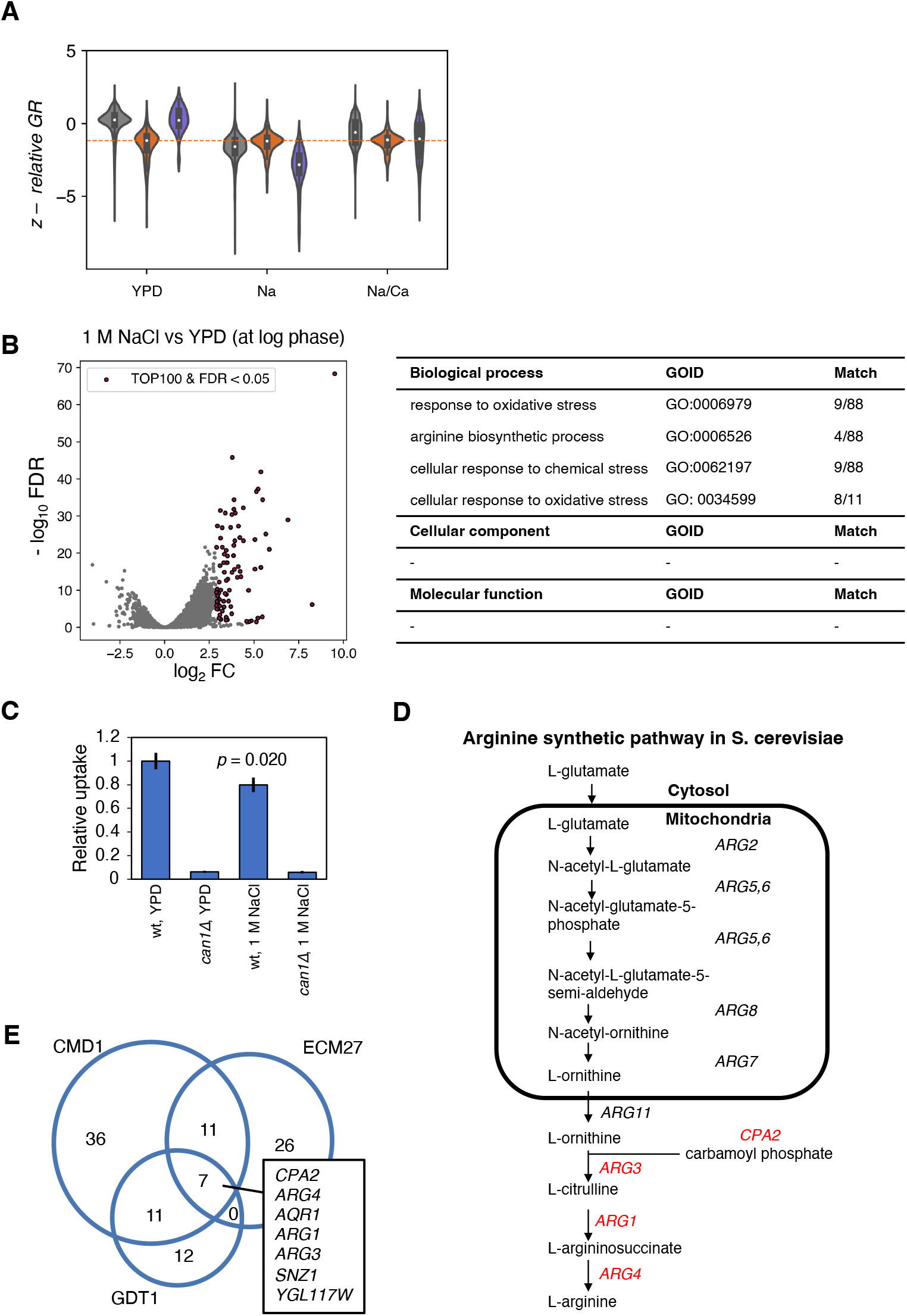
Transcriptome analysis of cells under salt stress obtained by RNAseq analysis. **A)** Relative fitness distribution when the growth rates in each condition were subtracted from Fig. 5F. **B)** Transcriptome changes in 1 M NaCl and YPD at log growth phase. ORFs with the top 100 most enormous fold changes and significant changes (*FDR* < 0.05) compared with YPD are highlighted. Gene ontology terms enriched in highlighted ORFs are shown in the left table. **C)** Arginine intake assay under YPD and 1 M NaCl. The *p*-values are from Welch’s t-test (n = 3). The error bars indicate SD. **D)** Scheme showing arginine synthesis pathway in *S. cerevisiae.* Red-colored genes were upregulated under 1 M NaCl and downregulated in GOFA’s overexpression stains. **E)** Venn diagram showing downregulated genes (*FDR* < 0.05) in GOFA’s overexpression stains. These data are summarized in Table S7.

**Extended Data Figure 10.**
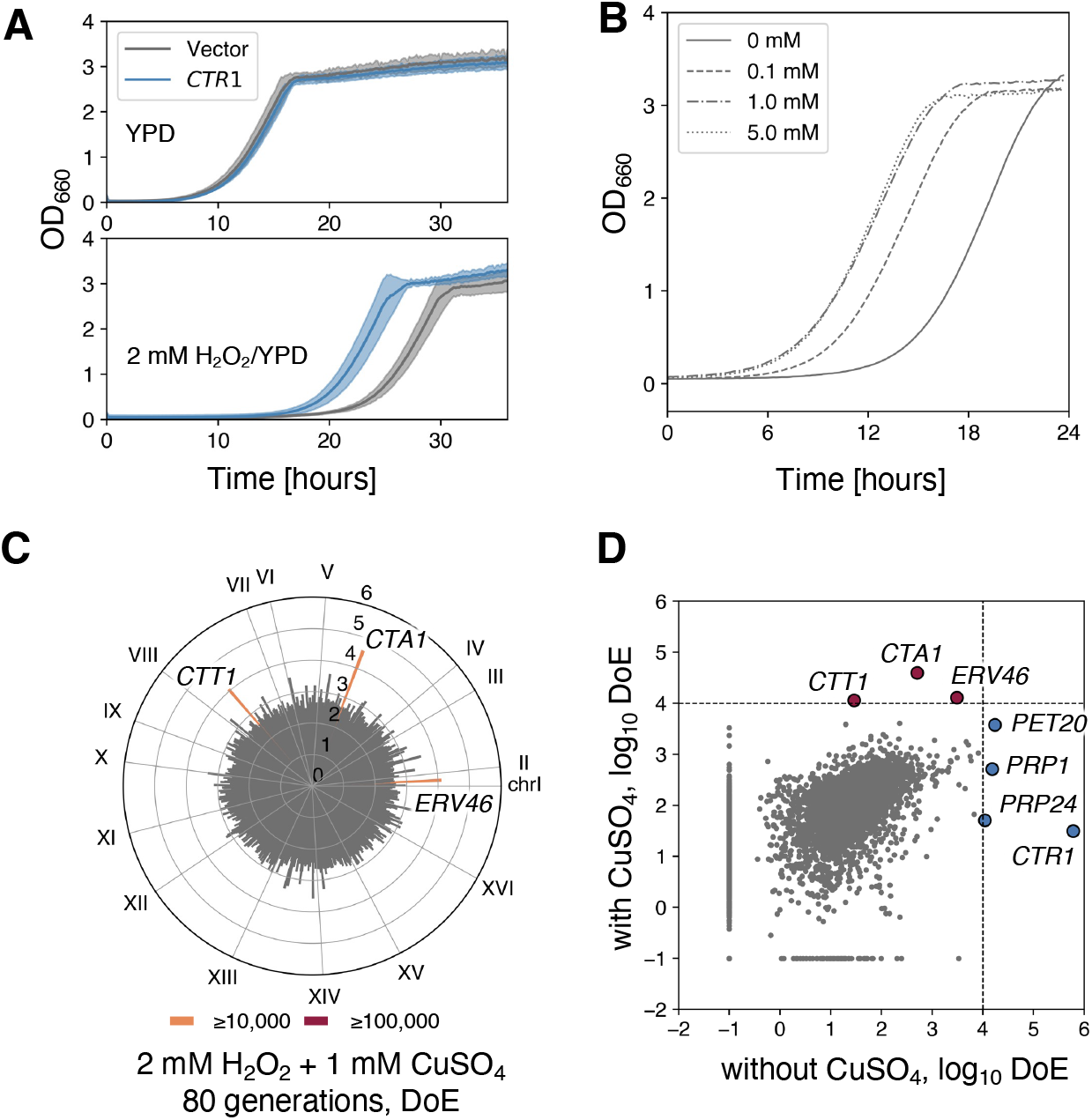
GOFAs enriched under oxidative stress propose Cu^2+^ limitation in the culture medium. **A)** CTR1-oe increased the growth of BY4741 under oxidative stress. Upper and lower panels indicate growth in YPD and 2 mM H_2_O_2_/YPD. The filled area shows SD (n = 3). **B)** The addition of CuSO_4_ increased the growth of BY4741 under 2 mM H_2_O_2_. **C)** DoE of plasmid inserts after the 80 generations-cultivation of ADOPT 1.0 library under 2 mM H_2_O_2_ with 1 mM CuSO_4_. The cycle chart shows the average DoE in four replicates. GOFAs are shown with red (DoE ≥ 100,000) and orange (DoE ≥ 10,000) bars, respectively. **D)** A comparison of the average DoE with and without CuSO_4_ addition. The colored circles indicate GOFAs, with (red) and without CuSO_4_ addition (blue). The dash lines represent the threshold of GOFAs as DoE ≥ 10,000.

